# Imaginal disc growth factors are *Drosophila* Chitinase-like Proteins with roles in morphogenesis and CO_2_ response

**DOI:** 10.1101/2022.06.29.498179

**Authors:** Anne Sustar, Liesl Strand, Sandra Zimmerman, Celeste Berg

**Affiliations:** Department of Genome Sciences, University of Washington, Seattle, Washington; Department of Developmental Biology, Stanford University, Stanford, California

## Abstract

Chitinase-like proteins (CLPs) are members of the family 18 glycosyl hydrolases, which include chitinases and the enzymatically inactive CLPs. A mutation in the enzyme’s catalytic site, conserved in vertebrates and invertebrates, allowed CLPs to evolve independently with functions that do not require chitinase activity. CLPs normally function during inflammatory responses, wound healing, and host defense, but when they persist at excessive levels at sites of chronic inflammation and in tissue-remodeling disorders, they correlate positively with disease progression and poor prognosis. However, little is known about their physiological function. *Drosophila melanogaster* has six CLPS, termed Imaginal disc growth factors (Idgfs), encoded by *Idgf1*, *Idgf2*, *Idgf3*, *Idgf4*, *Idgf5*, and *Idgf6*. In this study we developed tools to facilitate characterization of the physiological roles of the Idgfs by deleting each of the *Idgf* genes using the CRISPR/Cas9 system and assessing loss-of-function phenotypes. Using null lines, we showed that loss-of-function for all six Idgf proteins significantly lowers fertility and viability and compromises germ cell migration. We also showed that Idgfs play roles in epithelial morphogenesis, maintaining proper epithelial architecture and cell shape, regulating E-cadherin and cortical Actin, and protecting these tissues against CO_2_ exposure. Defining the normal molecular mechanisms of CLPS is key to understanding how deviations tip the balance from a physiological to a pathological state.

## Introduction

Chitinase-like proteins (CLPs) are secreted glycoproteins with properties of cytokines and growth factors (Guan *et al*. 2020; Kawamura *et al*. 1999; Kzhyshkowska *et al*. 2016; Lu *et al*. 2022). Normally, CLPs aid in inflammatory responses, wound healing, and defense against parasites, but they also have a dark side (Di Rosa *et al*. 2016; Mazur *et al*. 2021; Pinteac *et al*. 2021; Sutherland *et al*. 2014). Excess levels of CLPs are associated with diseases exhibiting chronically inflamed tissues, such as cancer, asthma, and arthritis, and there they promote the disease process rather than mitigate it (Di Rosa *et al*. 2016; Kzhyshkowska *et al*. 2016; Lee *et al*. 2011; Mazur *et al*. 2021; Pinteac *et al*. 2021; Qureshi *et al*. 2011).

The enzymatically active members of the family 18 glycosyl hydrolases hydrolyze the glycosidic bonds of chitin, a polysaccharide present in plant, fungal, bacterial, and animal species (Fuhrman *et al*. 1992; Miyashita and Fujii 1993; Samac *et al*. 1990; Tharanathan and Kittur 2003; Watanabe *et al*. 1993). CLPs, on the other hand, lack hydrolytic activity due to substitution of a key amino acid (glutamic acid) in the protein’s catalytic domain (e.g., with glutamine in *Drosophila* Idgfs and human CHI3L2, or with leucine in human CHI3L1) (Kirkpatrick *et al*. 1995; Recklies *et al*. 2002; Varela *et al*. 2002). This mutation was facilitated by duplication of ancestral chitinases, followed by subsequent mutations and further duplications (BUSSINK et al. 2007). Over evolutionary time, CLPs acquired new functions that do not require chitinase activity.

Extensive research in humans has identified four CLPs: chitinase 3 like 1 (CHI3L1), chitinase 3 like 2 (CHI3L2), oviductal glycoprotein 1 (OVGP1), and stabilin-interacting chitinase-like protein (SI-CLP). Most studies have focused on the expression patterns of CLPs and their clinical relevance as potential biomarkers of disease or as targets for therapy (Mazur *et al*. 2021; Pinteac *et al*. 2021). Analyses of the associated molecular mechanisms are limited to a few studies identifying binding partners, receptors, or signaling pathways (Areshkov *et al*. 2012; Francescone *et al*. 2011; Fusetti *et al*. 2003; Guan *et al*. 2020; He *et al*. 2013; Kzhyshkowska *et al*. 2004; Kzhyshkowska *et al*. 2006; Libreros *et al*. 2013; Low *et al*. 2015; Malik *et al*. 2015; Shao *et al*. 2009; Zhang *et al*. 2009), but these studies relied on *in vitro* cell culture, binding assays, immunohistochemistry, or biochemical methods rather than whole-animal models. To understand the molecular mechanisms of CLP function in physiological and pathological contexts, we need to analyze function *in vivo*, preferably in a model organism that would allow definitive functional tests.

The six *Drosophila* Idgfs are encoded by genes located at four sites in the genome: *Idgf1*, *Idgf2*, and *Idgf3* are clustered together on the left arm of chromosome 2 (2L), *Idgf4* lies on the X chromosome, while *Idgf5* and *Idgf6* reside ^~^ 1.7 Mb apart on the right arm of chromosome 2 (2R) (Figure 1A and Table S1) (Kawamura *et al*. 1999; Kirkpatrick *et al*. 1995; Zurovcova and Ayala 2002). Although now defined with a common nomenclature, the proteins were originally identified through different routes. Idgf6 (previously called DS47) was isolated from media from cultured Schneider line-2 (S2) cells, which have macrophage-like properties (Kirkpatrick *et al*. 1995). Idgf1, Idgf2, Idgf3, and Idgf4 were isolated from conditioned media from imaginal disc cell culture (Clone 8 cells) (Kawamura *et al*. 1999). Idgf5 was identified by computer search of the *Drosophila* genome (Zurovcova and Ayala 2002).

**Figure 1.**
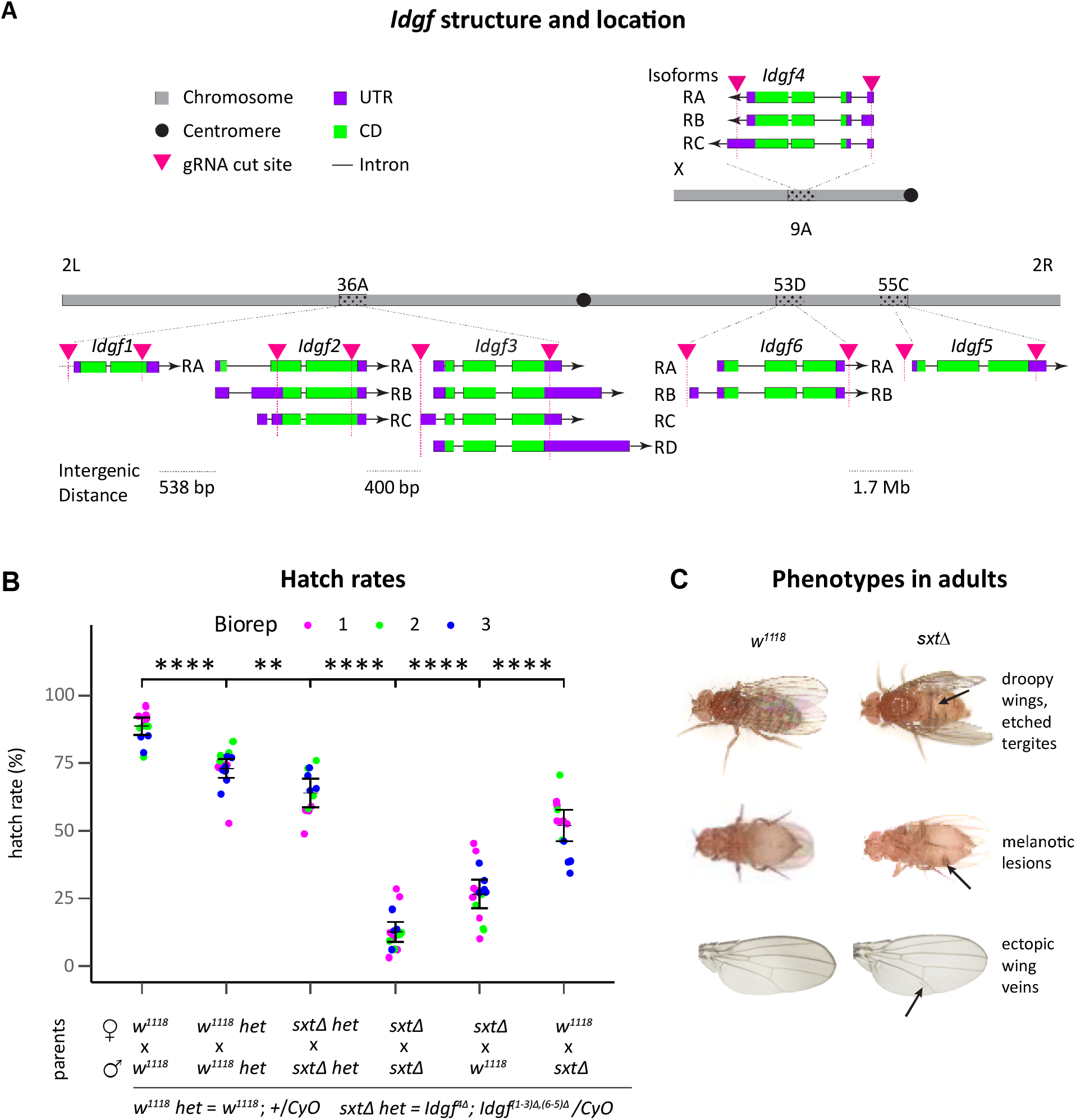
*Idgf* complete knockouts are not entirely lethal. (A) Diagram shows cytological locations of *Idgf* genes and structure of transcript isoforms, including 5’ and 3’ untranslated regions (purple), coding sequences (green), and introns (thin lines). Black arrows on transcripts indicate the orientation of transcription in the plus (right-pointing) or minus (left-pointing) direction. Arrowheads (magenta) indicate approximate cut sites for deleting each *Idgf* gene. (B) Homozygous and heterozygous *sxtΔ* mutants have low hatch rates relative to control (*w^1118^*). Dots represent egg laying assays sampled from three independent biological replicates, ranging from 350-852 eggs analyzed per replicate. Error bars indicate 95% confidence intervals. Significance is based pairwise t-tests with *p*-values adjusted for multiple comparisons (Benjamini-Hochberg), ** denotes *p*≤0.01, ****denotes *p*≤0.0001. (C) Adult phenotypes. Top row, dorsal view: etched tergites in adult abdomen (arrow). Wings are held out and down (“droopy” wings). Middle row, ventral view: dark cuticle patches (arrow) are reminiscent of melanotic clots. Bottom row: adult wings display ectopic wing veins (arrow). Anterior is up, distal is to the right.

*Idgf* proteins exhibit about 50% amino acid identity with each other and 15–25% amino acid sequence homology to family 18 glycosyl hydrolases, which include the chitinases (Kawamura *et al*. 1999; Varela *et al*. 2002). Consistent with their identification as secreted proteins, Idgfs and their human orthologs have an N-terminal signal sequence and an N-linked glycosylation sequence (Arias *et al*. 1994; Hu *et al*. 1996; Kawamura *et al*. 1999; Kirkpatrick *et al*. 1995; Meng *et al*. 2010; Nyirkos and Golds 1990; Renkema *et al*. 1998; Schimpl *et al*. 2012). The proteins form a characteristic barrel structure composed of eight parallel beta sheets surrounded by eight anti-parallel alpha helices (Varela *et al*. 2002).

Idgfs promote growth, proliferation, polarization, and motility in cultured cells (Kawamura *et al*. 1999). They participate in wound healing, protect against infection by nematodes (Broz *et al*. 2017; Kucerova *et al*. 2016), function in detoxification of cells (Broz *et al*. 2017), and participate in extracellular matrix organization required for molting (Pesch *et al*. 2016). To accomplish these functions, the Idgfs are not co-regulated, but rather, exhibit distinct expression patterns and respond to different stimuli. In the embryo, *Idgf* RNA is localized in cells adjacent to invaginating cells during tissue morphogenesis (Kawamura *et al*. 1999). In larvae and adults, Idgfs are expressed by hemocytes and the fat body (functional equivalent of the liver) and secreted into the hemolymph (Carton and Nappi 2001; Irving *et al*. 2001; Kawamura *et al*. 1999; Khush and Lemaitre 2000; Kirkpatrick *et al*. 1995; Meister *et al*. 2000). Hemocytes and fat body both participate in innate immune responses. In the ovary, Idgfs are expressed in dynamic patterns specific to the different cell types of the egg chamber (Zimmerman *et al*. 2017). Both over- and under-expression in egg chamber cells disrupt epithelial tube morphogenesis (Espinoza and Berg 2020; Zimmerman *et al*. 2017). Thus, Idgfs function in diverse processes in different tissues.

In this study, we focused on characterizing the physiological roles of the Idgfs by precisely deleting each of the Idgfs using the CRISPR/Cas9 system and assessing the effects of complete loss of function in various tissues and different life stages. We found that flies lacking all six Idgfs have low viability and fertility. Germ cell migration in these mutants is defective and likely contributes to the observed low fertility. We detected numerous cuticle defects in adults, abnormal epithelial morphogenesis in egg chambers, and segmentation defects in embryos. Finally, we found that Idgfs regulate E-cadherin and cortical Actin and protect epithelia against CO_2_ exposure.

## Results

### *Idgf* loss of function phenotypes

To begin to understand how Idgfs function, we developed *Idgf* null lines using the CRISPR/Cas9 system. We precisely deleted the coding region of each *Idgf*, creating single, double, triple, and more mutant lines, as well as a line with all six deletions (*w^1118^ Idgf^4Δ^*; *Idgf^(1-3)Δ,(6-5)Δ^*, termed “sextuple mutant” and designated from here on as *sxtΔ*) (Figure 1A, Figure S1, and Table S2). In the laboratory environment of rich food and benign growth conditions, we found no highly penetrant defects in any single mutant. *sxtΔ* flies, however, exhibited numerous phenotypes described below.

### sxtΔ mutants have low viability and fertility

Viability and fertility depend on many developmental processes that require patterning, growth, and cell migration. We assessed the function of Idgfs in this context by quantifying hatch rates for offspring from *sxtΔ* homozygous flies and from *sxtΔ* flies carrying the second chromosome CyO balancer (*w^1118^ Idgf^4Δ^*; *Idgf^(1-3)Δ,(6-5)Δ^/CyO*). We compared these rates to hatch rates of offspring from *w^1118^* and *w^1118^* carrying the same second-chromosome balancer (*w^1118^*; +/*CyO*) (Figure 1B).

The expected hatch rate for offspring from *w^1118^*; +/*CyO* parents is 75% of all laid eggs (*CyO/CyO* is lethal). In this background, we observed a hatch rate of 73% compared to a hatch rate of 63% in offspring from *w^1118^ Idgf^4Δ^*; *Idgf^(1-3)Δ,(6-5)Δ^/CyO* parents. The lower hatch rate for embryos in the *sxtΔ* background indicates that reduction of Idgfs moderately impacts embryonic development. For the heterozygous offspring from this cross (embryos carrying the *CyO* chromosome), maternal and zygotic contributions of wild-type Idgfs comes only from the *CyO* chromosome. In the *sxtΔ* homozygous embryos, all zygotic expression of Idgfs is lost. Consistent with maternal loading of *Idgf* transcripts (Zimmerman et al. 2017), maternal and zygotic loss of Idgfs is more detrimental: nearly 90% of eggs laid by *sxtΔ* females mated to *sxtΔ* males fail to hatch. Surprisingly, some escapers survive to adulthood, and when crossed to *w^1118^* flies, these escapers are weakly fertile (Figure 1B).

Heterozygous offspring from *sxtΔ* females crossed to *w^1118^* males have a higher hatch rate (26.5%) than embryos from a cross between *sxtΔ* homozygotes, presumably due to zygotic expression of wild-type Idgfs, and embryos from *w^1118^* females crossed to *sxtΔ* males, which have both maternal and potential zygotic *Idgf* expression in embryos, have a 52% hatch rate (Figure 1B). This percentage is lower than the hatch rate from the *Idgf^4Δ^*; *Idgf^(1-3)Δ,(6-5)Δ^/CyO* cross, possibly reflecting greater fertility of the male carrying the *CyO* balancer, which expresses wild-type Idgfs. These results indicate that both maternal and zygotic expression of wild-type Idgfs participate in embryonic development.

To further explore the effect of loss of Idgfs on viability and development, we compared eclosion rates between homozygous (straight-winged) and heterozygous (curly-winged) adult offspring of heterozygous parents (*w^1118^*; +/*CyO* or *w^1118^ Idgf^4Δ^*; *Idgf^(1-3)Δ,(6-5)Δ^/CyO* males and females). Considering the lethality of homozygous *CyO/CyO* flies, the expected Mendelian fractions of curly vs. straight-winged adults should be 2/3 heterozygous (curly-winged) and 1/3 homozyous (straight-winged) flies. Chi-squared analysis demonstrates that the relative fractions in the *sxtΔ* background (73% curly, 27% straight) deviates significantly from the relative fractions of wing phenotypes in the *w^1118^* background (69% curly, 31% straight) (*p* = 0.0087) (Figure S2A). Furthermore, *sxtΔ* homozygotes are developmentally delayed compared to *w^1118^* homozygotes. The fraction of *sxtΔ* homozygotes eclosing each day relative to the total after 3 days lags behind the *w^1118^* homozygotes (Figure S2B).

Taken together, these results indicate that loss of all six Idgfs impacts embryonic development, delays embryonic, larval and/or pupal development, reduces overall adult viability, and lowers both male and female fertility.

### Germ cell migration is defective in sxtΔ

In addition to low hatch rates, *sxtΔ* females laid fewer eggs than control females. Consistent with this observation, ovaries from *sxtΔ* adult females were greatly reduced in size compared to control females. Low fertility in *sxtΔ* flies may be the result of defects in germ cell migration and gonad development. Germ cells form at the posterior pole of the embryo and migrate through the gut, guided by attractive and repellent cues, to the somatic gonadal precursors; together these two cell types form the gonads (Bejsovec and Martinez Arias 1991; LeBlanc and Lehmann 2017). To assess whether low fertility was a consequence of defects in gonad formation, we marked germ cells with anti-Vasa antibody in control (*w^1118^*) and *sxtΔ* embryos (Figure S3A, A’). In the mutants, germ cell number is significantly reduced relative to embryos from control flies (Figure S3B, *p*=0.0002). These defects potentially involve guidance cues, cell adhesion, or polarization. In support of this hypothesis, we have observed a close association of *Idgf6::GFP* transgene expression with germ cells during embryonic Stages 4-7 (Fig S3C).

### Embryos exhibit segmentation defects

The insect body is organized in a series of segments, regulated by segmentation genes consisting of gap, pair-rule, and segment polarity genes. For example, *engrailed* is a segment polarity gene that is expressed in the posterior two rows of each segment (Figure 4A) and participates in the development of denticle belts (Glover and Hames 1989, pp.65-66). Denticles are hook-like cuticle structures on the ventral side of each larval segment that provide traction for crawling (Figure S4B). During late embryogenesis, the larval epithelial cells that secrete the ventral cuticle have one of two fates: either they remain smooth, or they produce polarized, actin-based protrusions that form the denticles (Dilks and DiNardo 2010) (Figure S4C). Maintenance of the segmental pattern of alternating smooth and denticle-producing stripes is regulated by the interactions between Engrailed (En), Wingless (Wg), and Patched (Ptc) (Glover and Hames 1989, pp. 65-66). The pattern of En expression was disrupted in *sxtΔ* mutant embryos, both on a tissue level and within each cell (Figure S4A’,B’,C’,D’). Smooth cuticle overlapped denticle belts, parts of neighboring belts sometimes fused together, single rows of denticles localized within bands of smooth cuticle, and En aberrantly localized to cell membranes (Figure S4D,D’) We explored these defects by examining the segmental pattern of denticle-producing and non-denticle-producing cell shapes in late-stage embryos and discovered an array of altered cell shapes and sizes (Figure 4C,C’). This phenotype could be due to patterning defects, or, alternatively, the cells may have achieved proper cell fate but cannot maintain proper positioning within a segment due to loss of cell-junction integrity. Cell polarity may also be affected.

**Figure 2.**
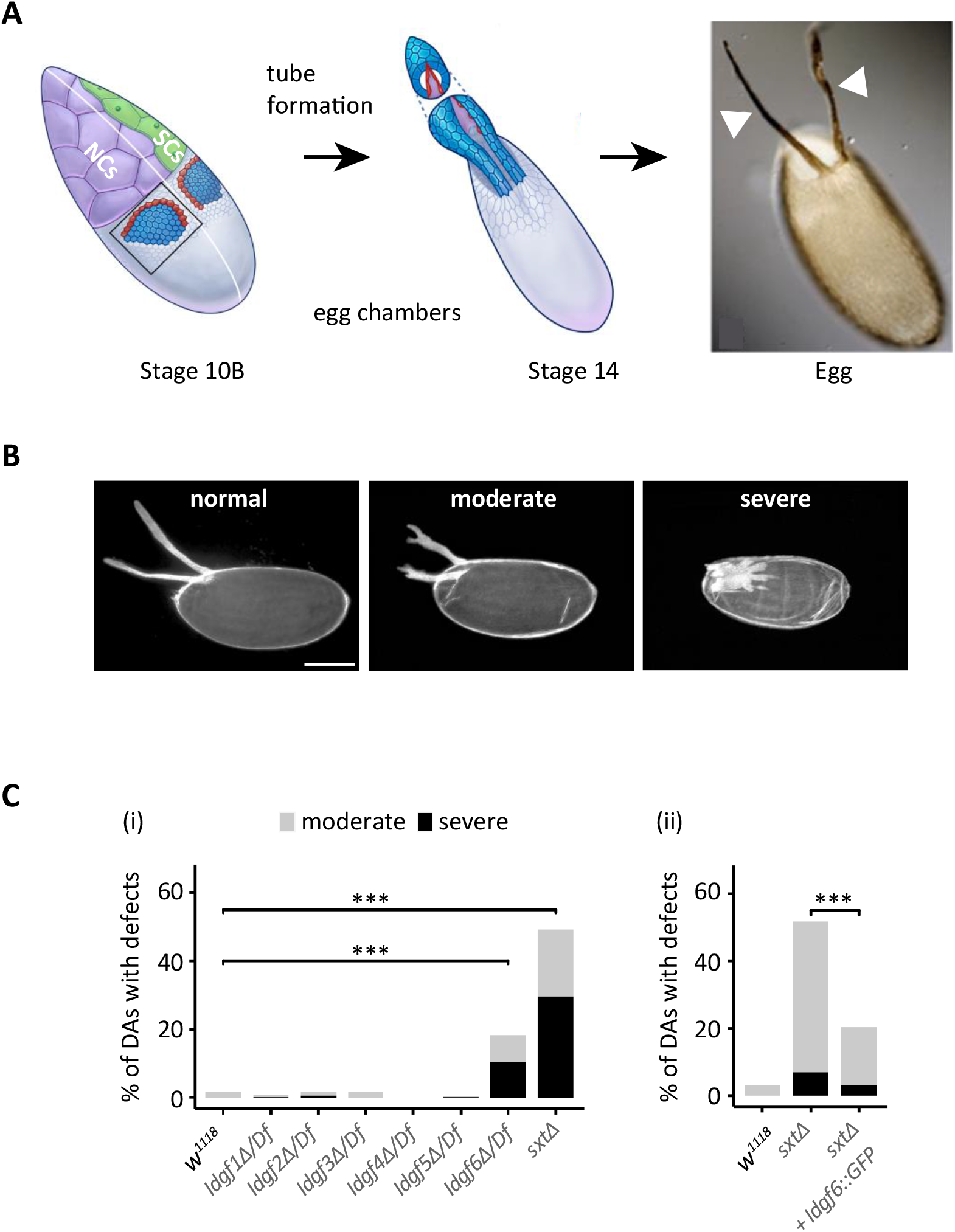
Dorsal appendage tube formation and defects in *Idgf* mutants. (A) Dorsal appendage formation. Roof cells (blue) and floor cells (red) change shape and reorganize to make two DA tubes. Stretch cells (green), cut away to show nurse cells (purple), guide tube elongation. Dorsal appendages (arrowheads) of the laid egg bring air to the embryo that is developing inside the eggshell. (B) Representative examples of DA morphology categorized into normal, moderate, and severe phenotypes. Scale bar = 100μm. (C) Moderate and severe DA phenotypes are significantly increased in eggs laid by *Idgf6*Δ/*Df* and *sxtΔ* females indicating defects in tubulogenesis during oogenesis. *Idgf1*Δ/*Df*, *Idgf2*Δ/*Df*, *Idgf3*Δ/*Df*, *Idgf4*Δ/*Df*, and *Idgf5*Δ/*Df* single mutants do not significantly affect DA morphology [C(i)]. Expression of a wild-type *Idgf6* transgene in *sxtΔ* mutant females partially rescues DA morphology [C(ii)]. Deficiency chromosomes specific to each mutant are listed in Table S3. *** indicates *p*<0.001, *X*^2^ test for trends in binomial proportions (Rosner 2000).

**Figure 3.**
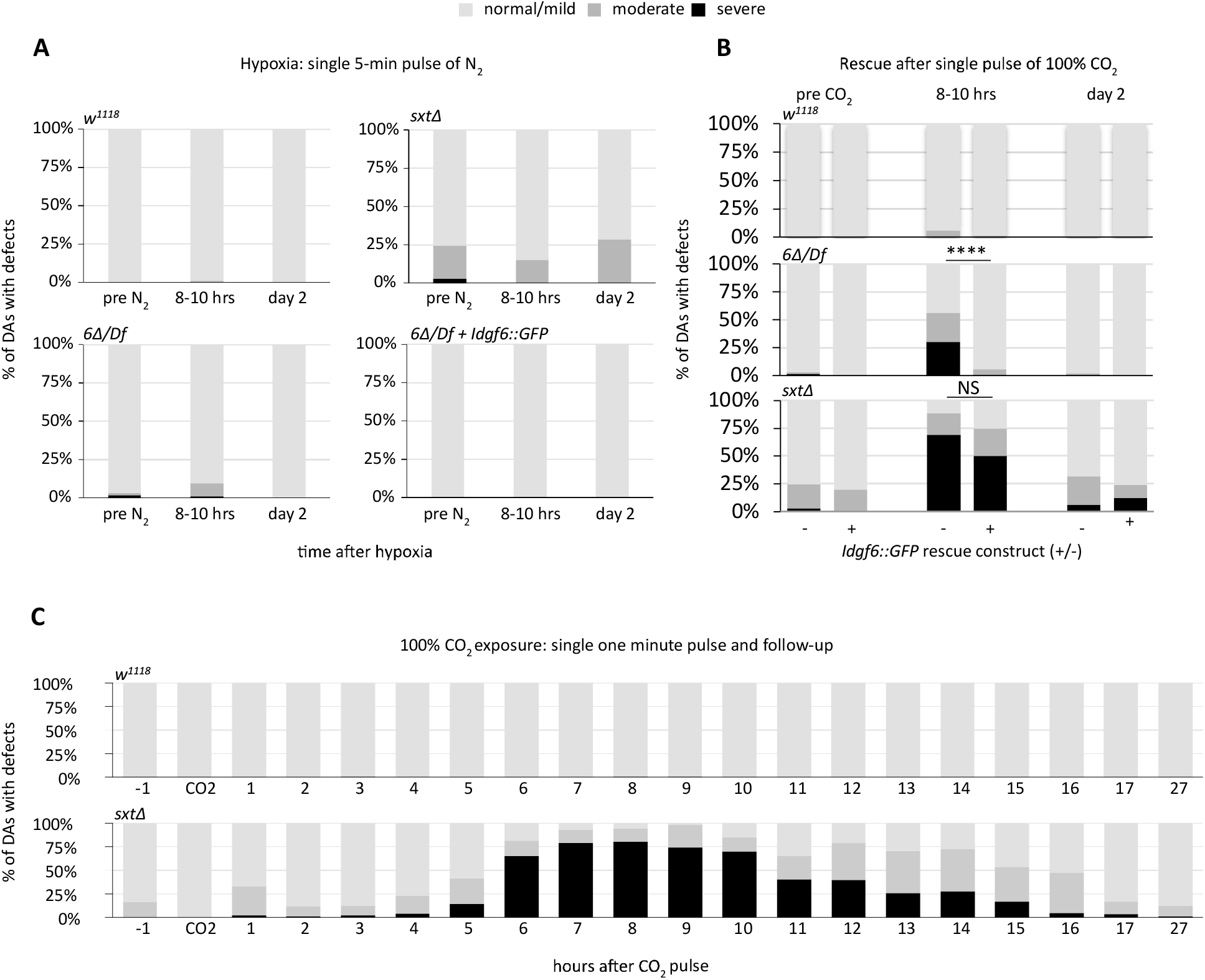
Idgfs protect against CO_2_ exposure. (A) Effect of hypoxia on DA phenotype. Air is replaced with 100% N_2_ for 5 min to induce hypoxia. DA phenotype is not significantly affected by hypoxia. (B) Expressing a single copy of a wild-type *Idgf6* transgene rescues the DA *Idgf6Δ* phenotype induced by a single, 1-min pulse of 100% CO_2_. The sxtΔ phenotype is partially but not significantly rescued (*p*=0.32). Proportions of defects peak at 8-10 hours after the CO_2_ pulse. The *Idgf6Δ* deletion is transheterozygous with a deficiency chromosome to cover potential background mutations. **** denotes significance (*p*≤0.0001, *X*^2^ test), NS=not significant. (C) Hourly assessment of DA phenotype following a 1-minute pulse of 100% CO_2_. DA defects in eggs laid by *sxtΔ* females peak at 7-10 hours after CO_2_ exposure.

**Figure 4.**
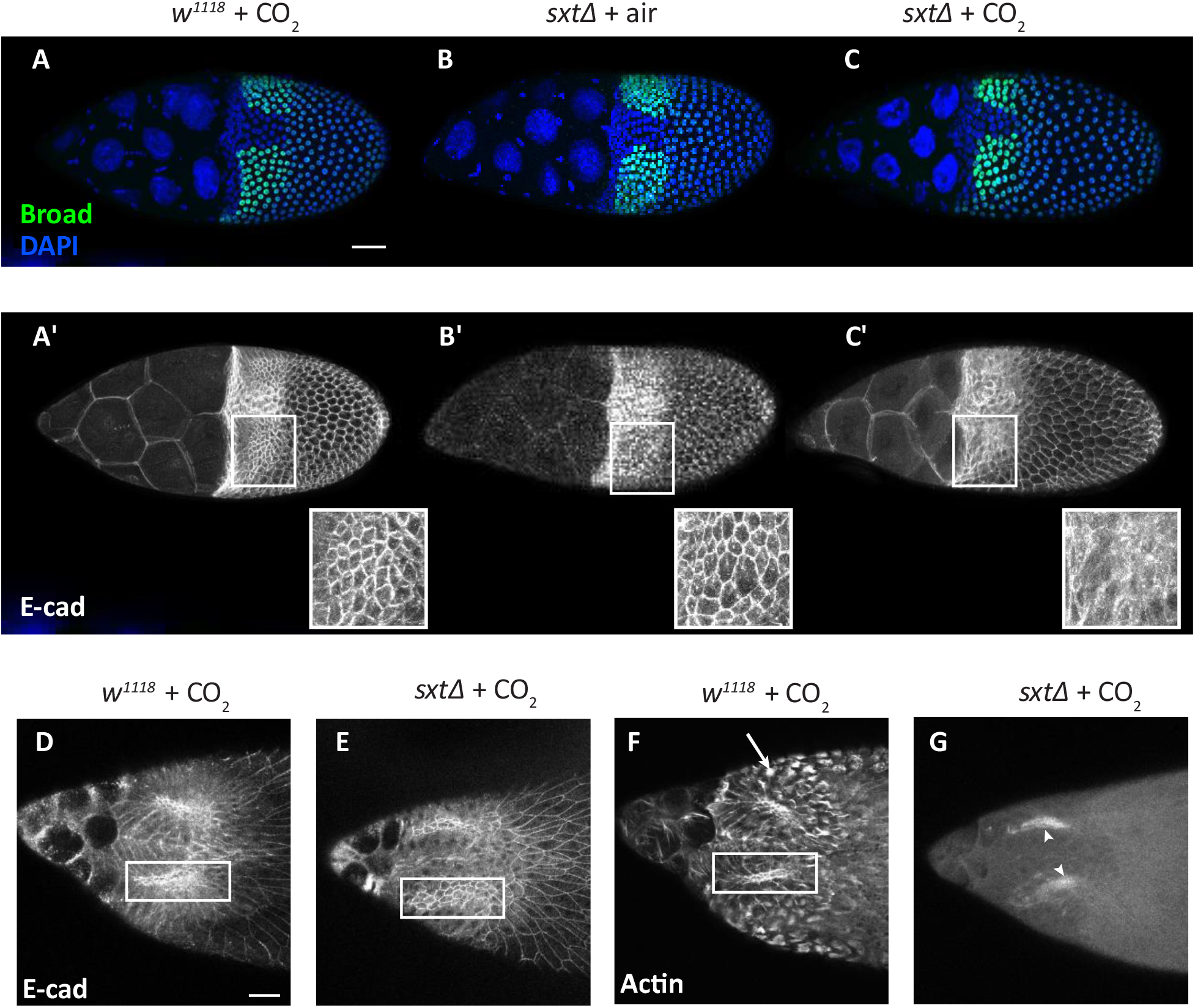
Patterning and cell morphology in DA-forming cells. (A-C’) Stage 10B egg chambers. Top row (A-C): Broad (green) marks the roof cells in the DA primordium, DAPI (blue) indicates the nuclei. Patterning is normal in *sxtΔ* mutant exposed to CO_2_. Lower row (A’-C’): E-cadherin delineates the apico-lateral cell junctions in follicle cells. Cell morphology is abnormal in *sxtΔ* mutant DAs, and E-cadherin localization is disrupted after a single pulse of 100% CO_2_ exposure (white rectangles indicate DA-forming cells). (D-G) Stage 12 egg chambers. DA roof cells (white rectangles) are abnormally large in the *sxtΔ* mutant (E) compared to control (*w^1118^*) (D) egg chambers. (F) As shown in *w^1118^* egg chambers, basal cell membranes normally exhibit bright spots of Actin localization (arrow). (G) In *sxtΔ* mutants, Actin is lost from cell membranes after CO_2_ exposure. Arrowheads in the *sxtΔ* + CO_2_ image indicate chorion autofluorescence.

### Adult flies display a variety of cuticle defects and lesions

Adult *sxtΔ* mutant flies display cuticle defects, i.e., etched tergites and “droopy” wings (Figure 1C, top row). Consistent with the role of Idgfs in immune responses (Kucerova *et al*. 2016), adults have lesions resembling melanotic clots, which normally occur at the sites of wounds or at sites of pathogen invasion (Figure 1C, middle row, arrow). Adult wings occasionally have ectopic wing veins (Figure 1C, bottom row, arrow), suggesting defects in wing imaginal-disk patterning during larval and early pupal development.

### Overall size of *Idgf* mutant adults was not affected

As noted in the Introduction, CLPs have properties of growth factors (Areshkov *et al*. 2012; Lee *et al*. 2011; Shao *et al*. 2009). To assess whether loss of Idgfs compromises growth, we quantified fly size by measuring wing area, which correlates well with overall body size (Mirth and Shingleton 2012; Siomava *et al*. 2016). We compared wing area in four different genotypes and found no significant difference under our culture conditions. Genotypes: *w^1118^*, *w^1118^* or *Idgf^(1-3)Δ^* in *trans* to a deficiency generated by the Bloomington Stock Center (NIH P40OD018537) (Ryder *et al*. 2004) that uncovers *Idgfs* 1-3 (Table S3), and *Idgf^(1-3)Δ^* in trans to a wild-type chromosome. We found no significant difference between any of these genotypes under our culture conditions. This result implies that these Idgfs do not contribute to growth and proliferation of imaginal discs and histoblast cells during larval and pupal development. Bristle morphology was also normal.

### Dorsal appendage tube formation is disrupted in sxtΔ mutants

Cell migration, polarization, growth, proliferation, and signaling govern the tissue movements and reconfigurations that form morphologic structures such as epithelial tubes. We used our dorsal appendage (DA) tube morphogenesis model to assess the effects of *Idgf* loss of function during oogenesis (Figure 2A). DA tubes produce eggshell structures that facilitate gas exchange for the developing embryo, and they are made by a subset of follicle cells in late-stage egg chambers. The egg chamber consists of 15 germline-derived nurse cells, one oocyte, and a surrounding monolayer of somatic epithelial cells consisting posteriorly of columnar cells and anteriorly of squamous cells (termed “stretch” cells). The tubes develop from two patches of precursor cells in the follicular epithelium and, through cell intercalation and cell shape change, elongate by migrating anteriorly over the stretch cells and beneath the extracellular matrix (Dorman *et al*. 2004; Osterfield *et al*. 2013; Rittenhouse and Berg 1995; Ward and Berg 2005). The tube cells secrete eggshell protein into the tube lumen, providing a readout for tube formation, similar to the way a mold forms a sculpture (DAs) (Figure 2A, right image). This highly conserved process of tube formation is a wrapping mechanism akin to mammalian neural tube formation (Osterfield *et al*. 2017).

We previously observed disruption of DA morphogenesis using *Idgf1* and *Idgf3* overexpression (Espinoza and Berg 2020; Zimmerman *et al*. 2017) and *Idgf* RNAi for each single mutant (Zimmerman *et al*. 2017). Since null mutations are the gold standard in genetics, we asked “What about complete knockout of *Idgfs*?” We compared the DA phenotypes of eggs laid by each single *Idgf* CRISPR-null mutant and the *sxtΔ* mutant to the DAs of eggs laid by control (*w^1118^*) flies and categorized the phenotypes as mild, moderate, or severe by scoring the DAs while blinded to the genotypes (Figure 2B). To limit the influence of potential background mutations, we assessed the phenotypes produced by single mutant genotypes *Idgf1Δ* through *Idgf6Δ* by placing each *Idgf* deletion chromosome *in trans* to a deficiency chromosome generated independently by the Bloomington Stock Center (NIH P40OD018537) (Ryder *et al*. 2004). Note that none of the deficiencies produced DA phenotypes on their own (Table S3). The *sxtΔ* mutant was assessed homozygously due to the unfeasibility of creating flies heterozygous with deficiencies spanning each of the six deleted *Idgf* loci.

Approximately 50% of eggs laid by the *sxtΔ* mutants and 18% of the *Idgf6*Δ mutants had moderate or severe phenotypes, but both of these percentages varied somewhat from experiment to experiment (see below). The other single mutants (*Idgf1*Δ, *Idgf2*Δ, *Idgf3*Δ, *Idgf4*Δ, and *Idgf5*Δ) did not exhibit significant levels of defects [Figure 2C (i)]. Expression of an *Idgf6::GFP* transgene in a *sxtΔ* mutant background partially rescued the *sxtΔ* DA phenotype [Figure 2C (ii)].

Overall, these results provide evidence for important physiologic roles of Idgfs in tissue morphogenesis, potentially including cell polarization, migration, adhesion, and signaling.

### Idgfs protect against effects of CO_2_ exposure

While analyzing the DA defects in eggs laid by *Idgf* mutants, we observed variability in penetrance and expressivity. Initially, we found differences in the percentage and severity of DA defects for eggs laid by *Idgf6Δ* females. Because these two sets of experiments (*Idgf6Δ* vs. other single *Idgf* mutants) were performed by two different lab workers, we asked whether these differences were biological or whether the variability was due to a difference in technique between lab workers (e.g., age of flies, overcrowding, or CO_2_ exposure). Intriguingly, we determined that there was variability in whether flies were exposed to CO_2_ when they were transferred to fresh egg-laying plates. This observation raised the question as to whether loss of *Idgf* function rendered flies more sensitive to CO_2_.

To compare CO_2_ sensitivity between wild-type and mutant flies, we assessed DA morphology on eggs laid by *w^1118^* flies and *Idgf* mutants by using several different regimes of CO_2_ exposure. Treating adult females to 100% CO_2_ for 60 seconds twice per day (^~^every 12 hours) for 3.5 days (Figure S5A) resulted in a significant increase in DA defects on the eggs laid by *Idgf6Δ* and *sxtΔ* mutants, but not on the *w^1118^* eggs nor in any of the other single mutants (*Idgf1Δ* through *Idgf5Δ*).

Since tissue and cellular responses to carbon dioxide depend on the length and severity of exposure (Helenius *et al*. 2016; Helenius *et al*. 2009; Sharabi *et al*. 2009a; Sharabi *et al*. 2009b), we asked if a prolonged exposure to a lower level of CO_2_ would have a similar effect. We exposed *sxtΔ* mutant and *w^1118^* adult flies to either 20% CO_2_ or 100% air and evaluated DA morphology at 1.5 – 2 days and 7-8 days of exposure. Exposure to CO_2_ over either of these time periods did not significantly increase DA defects in eggs laid by *sxtΔ* mutants (Figure S5B).

We next asked whether the effect of CO_2_ exposure was directly due to the CO_2_ itself or due to hypoxia, the transient absence of oxygen. To distinguish between these two hypotheses, we exposed flies to 100% N_2_ for five minutes followed by DA analysis at 8 – 10 hours and 2 days after exposure. We did not observe a significant effect on DA morphology following this treatment, indicating that the increase in DA morphology defects was directly due to CO_2_ exposure rather than hypoxia (Figure 3A).

These results suggest that Idgf proteins protect tissues against acute CO_2_ exposure. If this hypothesis is correct, expression of Idgf protein should rescue the DA mutant phenotype associated with high CO_2_. To test this prediction, we used a GFP-tagged transgene, *Idgf6::GFP*, in which Idgf6 is fused in-frame at its C-terminus to superfolder GFP (Sarov *et al*. 2016). We expressed this transgene in the *Idgf6Δ* and *sxtΔ* mutants and compared the DA phenotypes to the same mutants lacking the transgene. We exposed the flies to a single pulse of 100% CO_2_ for one minute and analyzed DA defects at 8 – 10 hours and 2 days after exposure (Figure 3B). While both mutants exhibited defects at 8 – 10 hours after exposure, *Idgf6Δ::GFP* expression restored the percentage of DA defects to pre-exposure levels in the *Idgf6Δ* mutants, and it reduced the DA defects in the *sxtΔ* mutants but not to the level of statistical significance. After 2 days, the DA defects in both mutants recovered to pre-exposure levels. These results demonstrate that this gene—environment interaction is due to the loss of Idgfs. Significantly, expression of a single Idgf protein can rescue or ameliorate the DA phenotype in *Idgf* mutants.

To examine the time course following acute CO_2_ exposure on a finer scale, we exposed adult *sxtΔ* flies to a single pulse of 100% CO_2_ for one minute, followed by hourly DA defect analysis (Figure 3C). Defects peaked at 6 – 10 hours after exposure and then recovered by ^~^27 hours after exposure.

In addition to the developmental roles for Idgfs in normoxia, these results support a hypothesis that Idgfs protect against morphogenetic disruption following acute CO_2_ exposure.

### Actin but not E-cadherin is disrupted in egg chambers after CO_2_ exposure

How do Idgfs function in DA morphogenesis, and how does acute exposure to CO_2_ impact that process? One hypothesis is that Idgfs are necessary for patterning the DA primordial cells. A second hypothesis is that Idgfs regulate cell morphology and epithelial tissue dynamics. To test these hypotheses, we exposed adult females to 100% CO_2_ for one minute and then waited for 20-30 minutes before dissecting, fixing, and staining the ovaries. We then examined patterning and cell morphology of the DA-forming cells using antibodies against Broad, a transcription factor that specifies the DA roof-forming cells, and against E-cadherin, an adhesion protein that outlines cell shapes (Dorman et al. 2004; Ward and Berg 2005; Ward et al. 2006).

Patterning was normal in both *w^1118^* and *sxtΔ* in both the exposed and unexposed egg chambers as revealed by two patches of ~ 50, high-Broad-expressing cells residing just lateral to the dorsal midline (Figure 4A-C). Cell morphology in the DA patches, however, was more disorganized, and E-cadherin was disrupted in the *sxtΔ* exposed egg chambers relative to the exposed *w^1118^* and unexposed *sxtΔ* egg chambers (Figure 4A’-C’, rectangles). DA cell circumferences were abnormally large in both the exposed and unexposed *sxtΔ* mutants relative to the DA cells in the exposed *w^1118^* egg chambers (Figure 4A’-C’, rectangles, and Figure 4D-E, rectangles). After exposure to CO_2_, *sxtΔ* DA-forming cells were further disrupted by a dramatic loss of Actin from both the cell junctions and the basal surfaces (Figure 4G). In contrast, cell shapes and Actin levels looked normal in the *w^1118^* DA-forming cells (Figure 4F).

We quantified E-cadherin and Actin in the DA-forming follicle cells by measuring fluorescent intensities across cell membranes using ImageJ (Figure 5, Figure S6) and calculating the total area under the plot profiles. To assess whether potential differences in intensity might be stage or cell-type specific, we analyzed roof cells, floor cells, and main-body follicle cells at Stage 10B, just prior to tube wrapping, and at Stage 12, during tube elongation (Figure 5A and Figure S6A). E-cadherin intensity was lower in *sxtΔ* mutants relative to *w^1118^* control flies (*p*≤0.05), but it was not significantly affected by CO_2_ exposure (Figure 5B,C and Figure S6B). In contrast, Actin levels were similar in *w^1118^* and *sxtΔ* mutants but dropped quickly in both samples upon exposure to CO_2_ (*p*≤0.05) (Figure 5B’,C’ and, Figure S6B’). After 30 minutes, Actin intensity recovered in the control egg chambers but remained low in the *sxtΔ* mutant samples (Figure 4F,G). These results suggest that Idgfs regulate E-cadherin expression or stability in normoxia and that they impact Actin dynamics in response to CO_2_.

**Figure 5.**
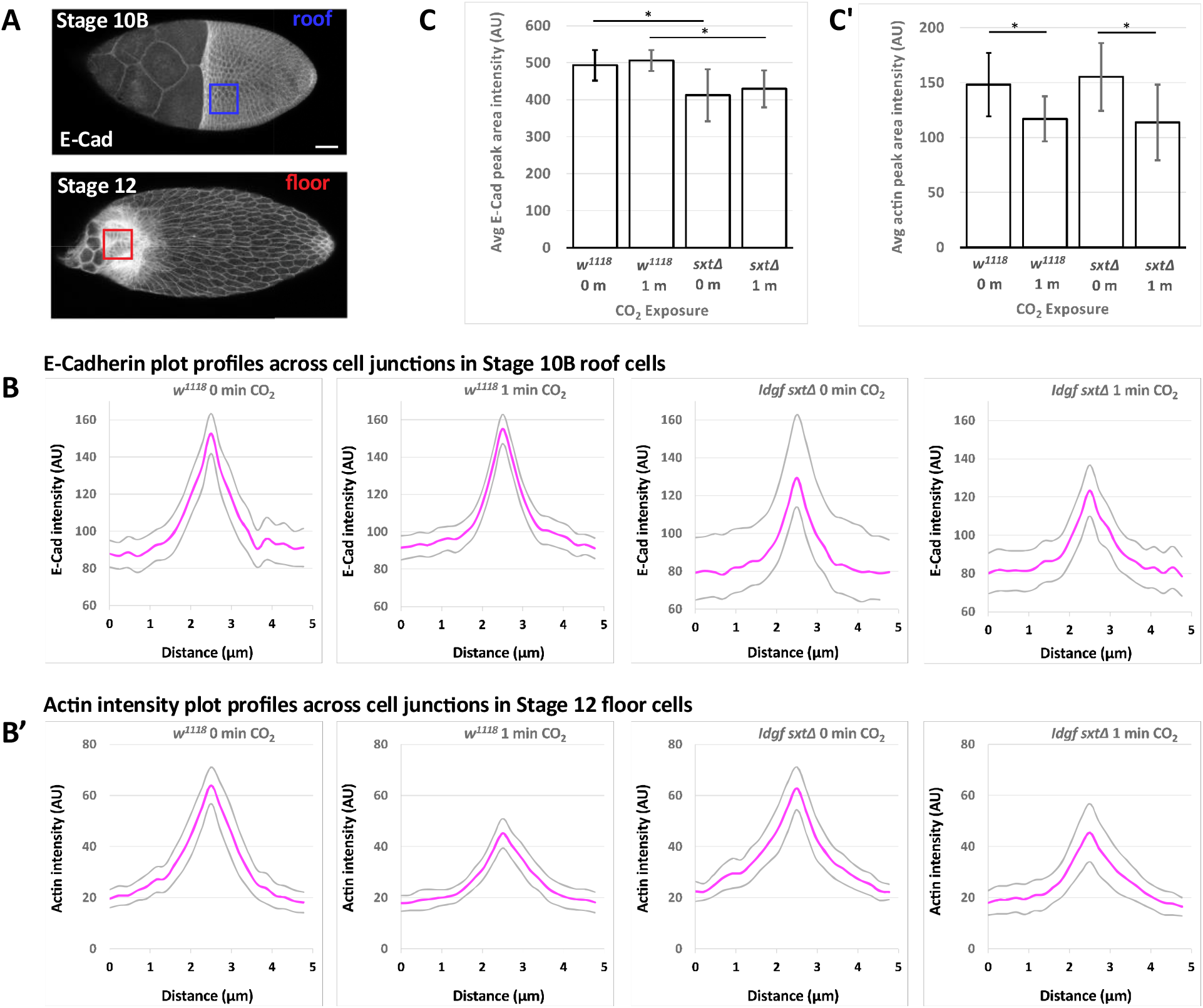
Quantification of cortical Actin and E-Cadherin intensity in DA cells. (A) E-Cadherin and Actin in Stage 10B roof cells (blue square) and Stage 12 floor cells (red square). Scale bar = 50 μm. (B) Comparison of *w^1118^* and *sxtΔ* fluorescence intensity plot profiles averaged across three cell membranes, from the center of one cell to the center of the neighboring cell, averaged over at least 6 egg chambers for each genotype and each exposure regime as indicated. Plots show mean (magenta) and confidence limits (gray). (C) Integrated area under the curves in B and B’ averaged over all measurements. Error bars show 95% confidence limits. * denotes *p*≤0.05, Student’s *t*-test, Benjamini-Hochberg adjusted *p*-values.

### Actin is disrupted in Stage 8-12 embryos after CO_2_ exposure

To ask whether Idgfs regulate E-cadherin and Actin in other contexts, we examined E-cadherin and Actin during embryonic development. We exposed *w^1118^* and *sxtΔ* embryos of all stages to 100% CO_2_ or pressurized air for 2 minutes before de-chorionating, fixing, and staining. Since embryos comprise numerous distinct cell types, we focused on cells near the surface that were relatively uniform in size and easy to measure. We chose ectodermal cells located between the mid-lateral cells and lateral edge of the germ band in abdominal segments 2-6 and measured these cells during the extended germ band stages (late Stage 8 – early Stage 12). Under these conditions, we observed no significant difference between E-cadherin levels in *w^1118^* embryos or *sxtΔ* mutant embryos under any of the three conditions (Figure 6A, Figure 6A’). Actin levels also did not differ significantly in *w^1118^* embryos between any of the three treatments (Figure 6B, Figure 6B’). However, Actin levels were significantly lower in *sxtΔ* mutants exposed to CO_2_ compared to unexposed embryos or embryos exposed to air in either genotype. Taken together, these results support our hypothesis that CO_2_ exposure disrupts Actin in embryonic epithelia.

**Figure 6.**
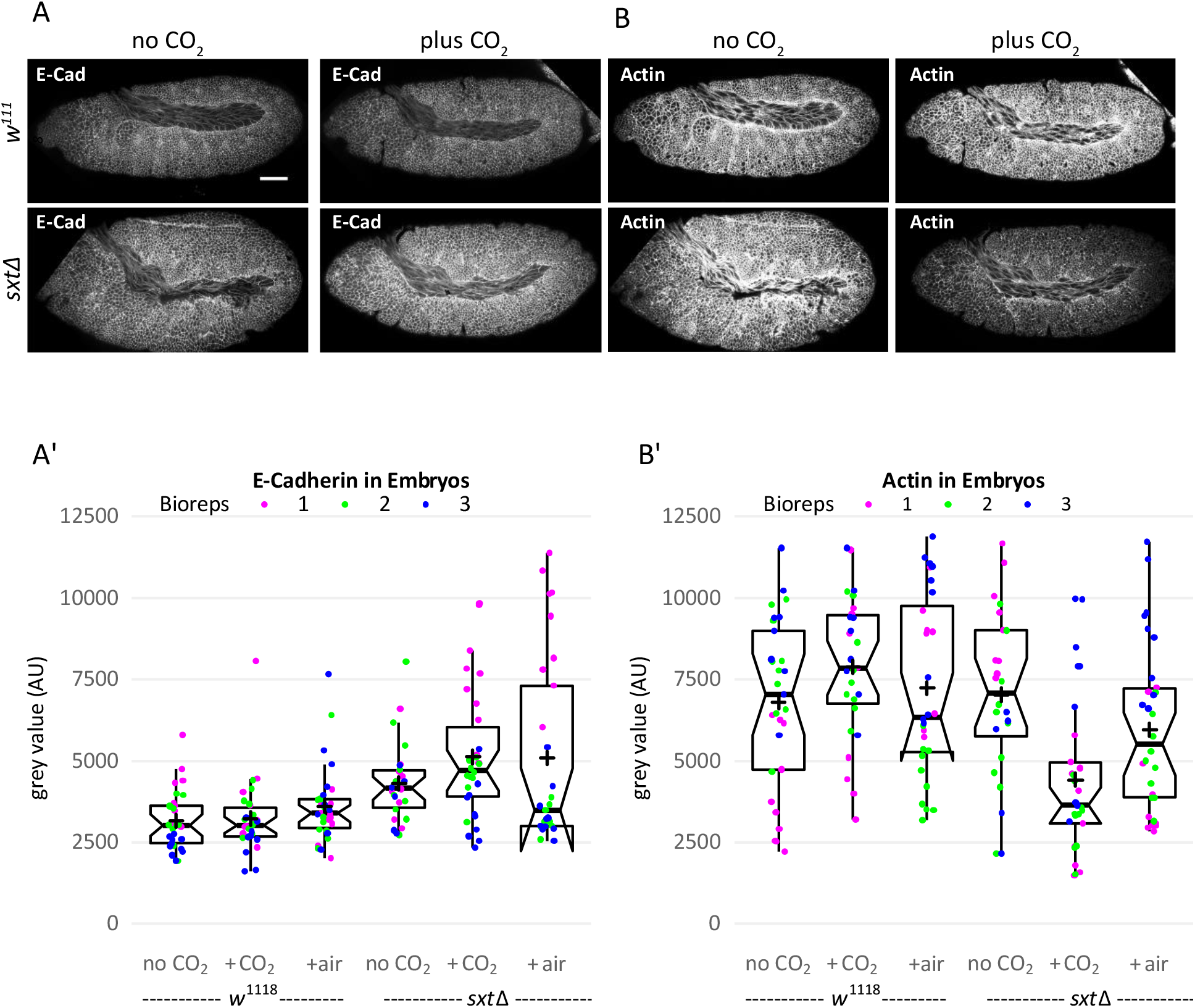
Quantification of cortical Actin and E-Cadherin intensity in Stage 8-12 embryos. (A, B) Representative embryo images showing E-cadherin (A) and Actin (Rhodamine phalloidin) (B) comparing genotypes and CO_2_ regime as indicated. Scale bar = 50 μm. (A’, B’) Comparison of *w^1118^* and *sxtΔ* fluorescence intensity measured across 6 cell membranes per embryo, from the center of one cell to the center of the neighboring cell. The colors represent different biological replicates. Each point in the graphs represents the average of the areas under the 6 plot profiles per embryo. The notches on the box plots represent the 95% confidence interval of the medians (horizontal lines), ±1.57*IQR/sqrt(n). IQR = interquartile range. + signs = means. Notches that double back occur when the uncertainty of the median exceeds the IQR. Non-overlapping notches are strong evidence that their medians significantly differ.

When analyzing Actin dynamics in egg chambers, we noticed that some cell types were more sensitive to CO_2_ exposure than others and that the effect of CO_2_ exposure on Actin perdured over a longer period in *sxtΔ* mutants compared to *w^1118^* controls. We therefore asked whether we might see more dramatic effects in embryos if we exposed embryos during cellularization, a period when Actin function is crucial (Sokac *et al*. 2022). We also reasoned that embryonic cells might be protected from CO_2_ exposure by the eggshell. We therefore exposed early-stage embryos to 100% CO_2_ and increased the exposure time to six minutes. Embryos were allowed to age for 30 minutes before fixing them and staining with phalloidin. We measured Actin fluorescence intensity across cell membranes in syncytial blastoderm embryos during stages 3-5. Prior to cellularization in Stage 5, nuclei in the syncytial blastoderm embryo are surrounded by Actin extensions (called pseudofurrows) at the periphery. In Stage 5, the Actin furrows extend around the nuclei to form a single layer of cells surrounding the yolky core. At these stages, the cells are uniform in size and shape, facilitating uniformity in measurements. As we observed in egg chambers, loss of Actin was significant in early embryos after exposure to CO_2_ but did not differ significantly between *w^1118^* and *sxtΔ* (Figure 7A,B,C).

**Figure 7.**
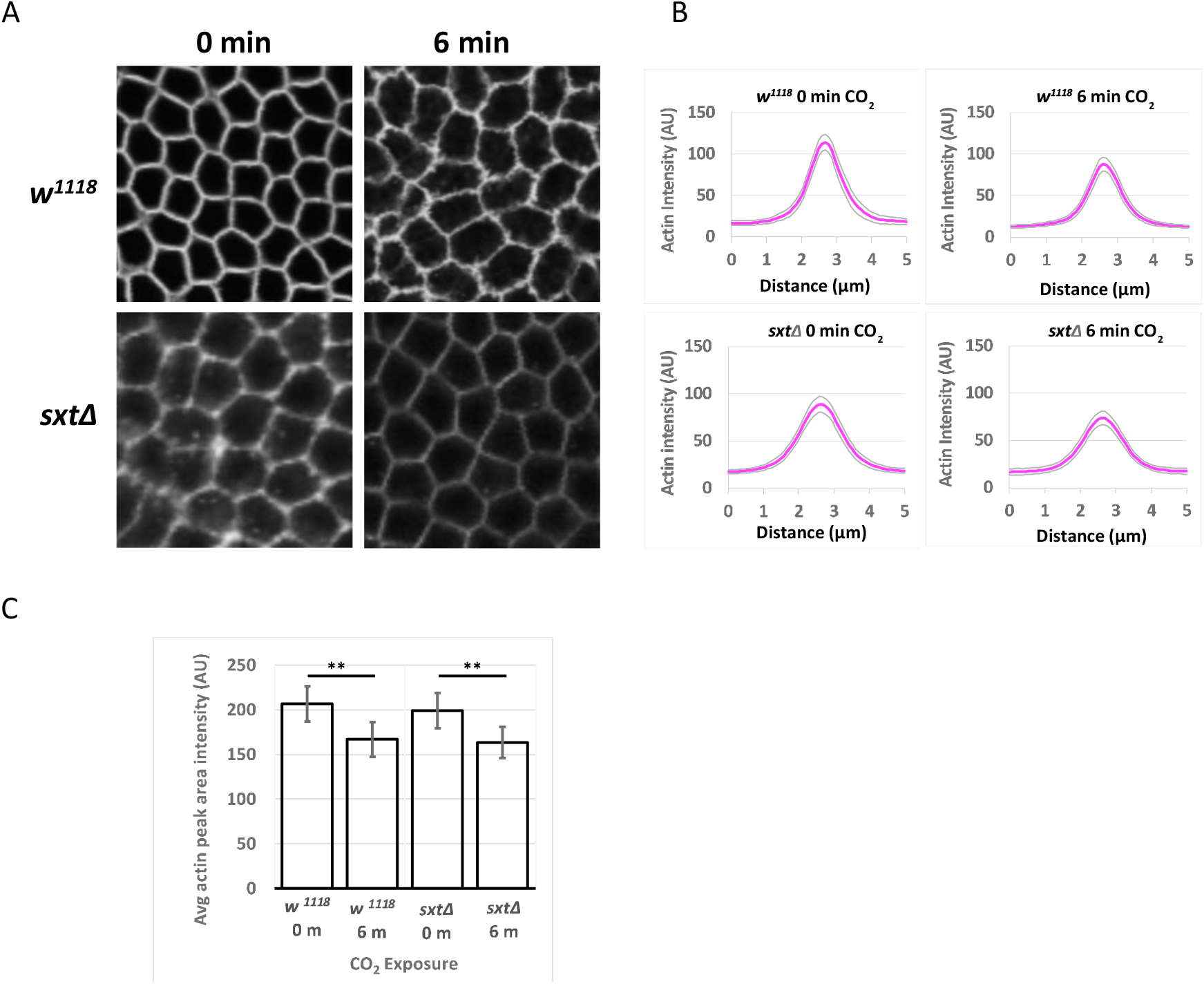
Quantification of cortical Actin intensity in Stage 3-5 embryos. (A) Actin in early embryos comparing genotypes and CO_2_ regime as indicated. (B) Comparison of *w^1118^* and *sxtΔ* fluorescence intensity plot profiles averaged across 6 cell membranes, from the center of one cell to the center of the neighboring cell, averaged over at least 10 Stage 3-5 embryos for each genotype and each exposure regime as indicated. Plots show mean (magenta) and confidence limits (gray). (C) Integrated area under the curves in B averaged over all measurements. Error bars show 95% confidence limits. ** denotes *p*≤0.01, Student’s *t*-test, Benjamini-Hochberg adjusted *p*-values.

## Discussion

We investigated the *in vivo* roles of *Drosophila* Imaginal disc growth factors throughout development. Previous studies have sought to characterize the function of individual Idgfs using RNAi or in cell culture. We are the first to delete each of the Idgfs and develop fly lines with single or multiple deletions of the Idgfs, including flies null for all six Idgfs. Using these tools, we found that loss of all Idgf function results in a variety of phenotypes.

In *Idgf sxt*Δ mutants, gonads form with significantly reduced germ cell numbers. Most germ cells are lost or die before they reach the somatic gonadal precursor cells, consistent with the low fertility we observed in *sxt*Δ adults. Normally, germ cells form at the anterior pole of the embryo and are taken into the midgut primordium during gastrulation. They initially form a tight cluster in the midgut, but subsequently extend protrusions toward the midgut cells, lose adhesion from each other, disperse, migrate individually through the midgut, and associate with somatic gonadal precursors cells to form the gonads (Review: Richardson and Lehmann 2010). To migrate, germ cells become polarized and extend protrusions toward the midgut cells. One hypothesis is that Idgfs participate in remodeling of cell junctions and Actin dynamics in migrating cells, consistent with our finding that loss of all Idgfs disrupts E-cadherin and Actin in certain contexts. A second hypothesis is that loss of Idgf function disrupts polarization and protrusiveness of primordial germ cells, consistent with a previous study demonstrating that Idgfs promote cell motility and protrusiveness in cell culture (Kawamura *et al*. 1999). Initial polarization of germ cells requires the G-protein-coupled receptor (GPCR) Trapped in endoderm 1 (Tre1), which is expressed in the germ cells and coincides with a redistribution of the Gβ subunit and adherens junction proteins such as Rho1 and E-cadherin (Kunwar *et al*. 2008). Tre1 is also important for migration across the midgut by signaling through small G proteins and the GTPase Rho1 (Kunwar *et al*. 2003). Studies have identified other key regulators of germ cell migration, such as the lipid phosphate phosphatases, Wunen and Wunen-2 (Hanyu-Nakamura *et al*. 2004), the small acidic protein, 14-3-3ε (Tsigkari *et al*. 2012), and pathways such as JAK/STAT (Brown *et al*. 2006; Li *et al*. 2003). CLPs are known to have properties of cytokines (Lee *et al*. 2011), raising the question of whether Idgfs may promote guidance cues for migrating germ cells. We have observed that Idgf6 associates with migrating germ cells from embryonic Stages 4-7; this association is lost by Stage 10. How Idgfs might interact with these components or otherwise participate in germ cell migration will require further study.

Consistent with studies showing expression of Idgfs throughout development, we showed that hatch rates are severely reduced in *sxtΔ* homozygous embryos from *sxtΔ* parents. This effect of loss of all maternal and zygotic Idgfs was ameliorated by adding either maternal, zygotic, or both maternal and zygotically expressed Idgfs. Interestingly, offspring of parents where both parents carry a *CyO* chromosome in a *sxtΔ* background have a higher hatch rate than offspring of *w^1118^* females crossed to *sxtΔ* males, even though both scenarios produce embryos that presumably receive both maternal and zygotically expressed Idgfs. Perhaps *sxtΔ* males are less fertile than their *sxtΔ* counterparts carrying the *CyO* chromosome, which expresses wild-type Idgfs. One hypothesis is that the sxtΔ male’s fertility is compromised by defective germ cell migration, consistent with our finding that germ cell migration is defective in *sxtΔ* embryos.

Further, we found that the proportions of homozygous vs. heterozygous offspring produced by heterozygous parents deviate significantly from the expected Mendelian ratios, reflecting greater lethality of the homozygotes from embryonic through adult stages. Our observation that eclosion is delayed in *sxtΔ* mutants could imply that Idgfs function as growth hormones; loss of function could slow development. Nevertheless, we found no significant size difference between *Idgf* mutants and controls.

Consistent with disruption of embryonic development (as revealed by lower *sxtΔ* hatch rates), we observed abnormal expression of Engrailed protein and disorganized denticle belts in embryos, including a mislocalization of Engrailed at the cell membrane. Patterning is achieved through the expression of the segment polarity genes *wingless* (*wg*) and *engrailed* (*en*), which activates expression of *hedgehog* (*hh*). Within each segment, cells receiving the Wg signal produce smooth cuticle posteriorly, and cells receiving the Hh signal produce denticles anteriorly (Bejsovec and Martinez Arias 1991; Swarup and Verheyen 2012). The denticle-versus-smooth cell shapes are distinct: whereas the denticle-producing cells are rectangular and arranged in rows with the long edges oriented in the ventrolateral direction (similar to a staggered brick-wall arrangement), the smooth cells are larger and not arranged in rows (Hirano *et al*. 2009). Mislocalization or abnormal expression of En could cause a transformation of cell fate from non-denticle-producing to denticle-producing cells, resulting in the disorganized denticle belt phenotype we observed in *sxtΔ* embryos (Figure S4B). Alternatively, cells in the epidermal epithelium may have altered cell adhesion and increased motility, allowing denticle-producing cells to invade into the non-denticle-producing cells.

Segmentation defects were apparent in adults in that *sxtΔ* flies exhibited etched tergites. Each abdominal segment in the adult epidermis develops from three bilateral pairs of groups of cells called histoblast nests. The anterior pair in each segment forms the tergite. Histoblast nests proliferate after pupariation and migrate by intercalation into the larval epithelial cells, which then extrude basally and die. The histoblast nests fuse and secrete the adult cuticle (Athilingam *et al*. 2021; Davis *et al*. 2022; Mangione and Martin-Blanco 2018; Michel and Dahmann 2020; Nardi *et al*. 2018; Panzade and Matis 2021; Prat-Rojo *et al*. 2020). Successful migration depends on Decapentaplegic (Dpp) and protrusive extensions at the leading edge of the histoblast nests (Ninov *et al*. 2007; Ninov *et al*. 2010). Our findings suggest a role for Idgfs in cell migration, but how Idgfs function in tergite formation is unknown.

Our previous studies showed that the stretch cells relay signals from the nurse cells to the DA cells to guide their movement and ensure tube closure. These signals rely on function of the SOX transcription factor Bullwinkle (Bwk) in the nurse cells [Rittenhouse dissertation, 1996. Bullwinkle, an HMG box protein, is required for proper development during oogenesis, embryogenesis and metamorphosis in *Drosophila melanogaster*, PhD, University of Washington; (Rittenhouse and Berg 1995)], and the non-receptor tyrosine kinases Shark and Src42a in the stretch cells (Tran and Berg 2003). The specific signaling molecules and their targets are unknown. *Bwk* mutants produce short, wide, open tubes that create DAs resembling moose antlers (Dorman *et al*. 2004; Rittenhouse and Berg 1995). Shark and Src42A act downstream of Bwk to regulate DA morphogenesis; Shark loss of function enhances the Bwk phenotype (Tran and Berg 2003).

In this study, we found that except for *Idgf6*, the single *Idgf-null* mutants did not cause a defective DA phenotype. The results for the single null mutants contrast with our previous studies using RNAi to knock down each *Idgf* individually (Zimmerman *et al*. 2017); we found that RNAi knockdown of each *Idgf* results in a partially penetrant DA phenotype featuring shorter, wider DAs. Shorter, wider DAs also occur when Idgfs are overexpressed or elevated as in eggs from *bwk* mutant flies and contrast with the moderate category of defective DAs in the null mutants, which have thin DAs. Why do significant defects occur in single *Idgf* RNAi knockdowns, but not in single nulls? Functional redundancy from the remaining Idgfs or even from non-Idgf genes could account for the incomplete penetrance or lack of a phenotype. Duplicated genes can maintain functional redundancy despite their divergence over evolutionary time (Dean *et al*. 2008; DeLuna *et al*. 2008; Kafri *et al*. 2008; Musso *et al*. 2008). Functional redundancy between paralogous genes can result in a strong phenotype when all of the paralogs are deleted and a weak or neutral phenotype when the genes are deleted singly (Dean *et al*. 2008; DeLuna *et al*. 2008; Gu *et al*. 2003; Kafri *et al*. 2008; Musso *et al*. 2008), similar to what we observed for the *Idgf* null mutants.

Compensation in single null mutants from the remaining wild-type paralogs or even from *non-Idgf* genes could be post-transcriptional, e.g., through increased translation or greater protein stability. Compensation may also occur through transcriptional adaptation, in which upregulation of compensating genes can be triggered by any of a number of mRNA decay pathways that target aberrant mRNAs (Garneau *et al*. 2007). A recent study demonstrated that transcriptional adaptation depends on the existence of a mutant mRNA, for which degradation can occur via different surveillance pathways, including non-stop, no-go, or nonsense-mediated decay, depending on the nature of the mutation. Upregulation can then occur in genes with sequence similarity to the degraded mRNA (El-Brolosy *et al*. 2019; El-Brolosy and Stainier 2017; Garneau *et al*. 2007; Rossi *et al*. 2015).

Except for *Idgf2Δ*, the single null mutants cannot express a transcript. The *Idgf2* deletion leaves intact the 5’ UTR, the first exon and intron, and part of the second exon, potentially allowing production of a mutant transcript and therefore transcriptional adaptation in the *Idgf2Δ* and *sxtΔ* mutants, which carry the *Idgf2* deletion. Degradation of the *Idgf2Δ* transcript could not be via the nonsense-mediated decay (NMD) process, which requires a premature stop codon upstream of an intron. Degradation via a non-NMD pathway, however, could be occurring in either the *Idgf^2Δ^* or *sxtΔ* mutant lines. Our RNAi experiments were performed in a wild-type background, so there would be no mutant mRNAs, and any adaptation would be on the translational or post-translational protein level (DeLuna *et al*. 2010; Donnelly and Storchova 2014; Ishikawa *et al*. 2017; Torres *et al*. 2008).

Although adaptation could be occurring in both the nulls and the knockdowns, the time frame for adaptation could also be a factor. In the RNAi study, we used the GAL4/UAS system to knockdown transcripts from each of the *Idgf* genes. We used a stretch-cell-specific Gal4 driver, which is not expressed until Stage 10 of egg chamber development, after patterning has occurred but just hours before tube formation. This timing may not allow enough time for compensation to occur. On the other hand, *Idgf-null* mutant flies are mutant throughout their entire life cycle. Thus, the different results of our *Idgf* RNAi knockdown and null experiments could be due to different mechanisms of compensation. Discerning whether compensation is occurring and what contributes to the differences in phenotypes exhibited by overexpression, RNAi, and gene deletion will require further study.

Mutations can render cells less robust to endogenous or exogenous perturbations (Levy and Siegal 2008; Masel and Siegal 2009). *Drosophila* researchers commonly use CO_2_ to anesthetize flies without considering the potential profound and long-lasting effects of CO_2_ exposure. Previous studies have shown that hypercapnia can suppress the immune system and alter gene expression via the NF-κB pathway in mammalian cells (Cummins *et al*. 2010). Hypercapnia also reduces fertility (Helenius *et al*. 2009) and alters climbing and flight behavior in *Drosophila* (Bartholomew *et al*. 2015).

We demonstrated that exposing *Idgf*-null mutants to CO_2_ enhances dorsal appendage morphogenesis defects independently of hypoxia and induces loss of cortical Actin in epithelial tissues during oogenesis and embryonic development. We did note molecular differences, raising the question of whether Idgfs regulate E-cadherin in normoxia and Actin in response to CO_2_. We saw that E-cadherin was disrupted in *sxtΔ* mutant egg chambers but was unaffected by CO_2_ exposure in either egg chambers or embryos. In contrast, Actin levels were unaffected in *sxtΔ* mutants vs. controls but significantly reduced in *sxtΔ* mutants upon CO_2_ exposure. It could be that the Actin is also perturbed in wild-type tissue but recovers, whereas absence of Idgfs eliminates a similar recovery in *sxtΔ* tissue. The mechanisms leading to these novel outcomes are unknown.

One possible mechanism for this regulation is that Idgfs interact with the immune deficiency (IMD) pathway through the transcription factor Relish (Drosophila NF-κB). Previous studies show that Relish inhibits JNK signaling (Park et al., 2004). Activation of JNK signaling reorganizes the Actin cytoskeleton, impacting developmental processes such as cell migration and dorsal closure (Homsy *et al*. 2006; Jacinto *et al*. 2000; Kaltschmidt *et al*. 2002; Kockel *et al*. 2001; Rudrapatna *et al*. 2014). Upon Relish loss, JNK activation causes upregulation of actin remodelers (Ramesh *et al*. 2021). Another possibility is that CO_2_ exposure reduces pH at the cell membrane, thereby altering Actin dynamics. pH is an important regulator in developmental processes such as differentiation (Ulmschneider *et al*. 2016) and tissue architecture (Grillo-Hill *et al*. 2015). Alternatively, the mechanism could involve a pathway independent of JNK signaling or pH.

A common thread among the various phenotypes produced by loss of all Idgfs is the disruption of E-cadherin and Actin, an outcome that could interfere with cell migration, segmentation, and tissue morphogenesis. Consistent with this hypothesis, we recently identified eukaryotic translation initiation factor 3 subunit e (eIF3e) as a strong enhancer of Idgf3 overexpression (Espinoza and Berg 2020). eIF3e is part of a complex that regulates the redox enzyme, Mical, which functions to disassemble Actin filaments and inhibit Actin polymerization (Grintsevich *et al*. 2021; Grintsevich *et al*. 2017; Grintsevich *et al*. 2016; Hung *et al*. 2010). We are currently beginning to investigate the mechanistic role of Idgfs in this pathway.

Our investigation into the effects of complete loss of function of Idgfs has raised intriguing questions regarding Idgf function. What are the pathways? What are the precise mechanisms? What are the common threads among the various phenotypes? Our *Idgf* null lines are valuable tools that will facilitate studies providing new insights into the function of Idgfs in *Drosophila*. Our results suggest new avenues of investigation for the mechanisms of CLP function in cancer pathogenesis, autoimmune disease, and tissue remodeling disorders.

## Materials and Methods

### Fly stocks

Bloomington stocks: *w^1118^* (3605), *Df(2L)ED1102* (24113), *Df(1)ED6989* (9056), *Df(2R)BSC337* (24361), Df(2R)Exel6064 (7546). VDRC stock: Idgf6^fTRG01331.sfGFP-TVPTBF^ (Sarov *et al*. 2016). Stocks generated in this study: *Idgf^1Δ^*, *Idgf^2Δ^*, *Idgf^3Δ^*, *Idgf^4Δ^*, *Idgf^5Δ^*, *Idgf^6Δ^*, and Idgf^4Δ^; Idgfp ^(1-3)Δ,(6-5)Δ^ (*sxtΔ*) were used in the experiments. All *Idgf* mutants generated in this study, including double and triple nulls, are listed in Table S2. Flies were maintained on standard molasses food at 25°C.

#### Reagents and materials

<u>Ovaries</u>

- Dissecting dishes and forceps
- PBS, 10x (Sambrook et al. 1989), NaCl 80 g, KCl 2 g, Na2HPO4 14.4 g, KH2PO4 2.4 g in 1 L of H2O pH 7.4.
- PBT, 0.1% (vol/vol) Tween 20 diluted in 1× PBS
- EBR, 10× (modified Ephrussi-Beadle Ringer’s solution) 1.3 M NaCl, 47 mM KCl, 19 mM CaCl2 and 100 mM HEPES (pH 6.9).
- Paraformaldehyde, 4% (wt/vol), Dilute 16% (wt/vol) paraformaldehyde in 1× PBS at a ratio of 1:4
- Western Blocking Reagent (WBR)

<u>Embryos</u>

- Flow Buddy - Benchtop Flow Regulator (Genesee Cat #: 59-122B)
- Egg collection basket - Falcon tube (50 mL) with the bottom cut off and nylon mesh attached to bottom (e.g., melt the bottom edge slightly with a Bunsen burner and press on the nylon mesh)
- Double-sided sticky tape
- Whatman paper
- Pipette controller (e.g., Millipore Sigma BR25900)
- Pasteur pipettes, glass, 3.5 inches
- Petri dish lid
- Plastic fly bottles punched with holes (to make egg laying chambers when attached to apple juice plates)
- 30G syringe needles + plastic syringes
- PBTA with 0.05% or 0.1% Triton (1x PBS, 1% BSA, 0.05% or 0.1% Triton X-100, 0.02% Sodium Azide)
- Apple juice plates, 175 mL H2O, 25 mL 4x apple juice, 5 g of agar 1.2 g of methyl paraben. Heat in microwave until agar dissolves. Pour into Petri dish lids. These will be attached to the plastic fly bottles using rubber bands.
- Bleach
- Heptane Saturated with 37% Formaldehyde - Combine equal volumes of heptane and 37% formaldehyde, shake the mixture vigorously for 15 seconds. Let the solution settle into the 2 phases. Prepare the solution the day before it is to be used, shaking the vial or bottle periodically throughout the day. The saturated heptane is the upper phase in the 1:1 heptane:formaldehyde stock. Store at room temperature, protected from light, up to several months.
- 1x Embryo Wash Solution, - 0.7% NaCl, 0.05% Triton X-100, H_2_O
- Normal goat serum (NGS)

#### Tissue processing and Immunohistochemistry

Ovaries - Ovaries were dissected in EBR, fixed for 20 minutes in 4% (wt/vol) paraformaldehyde, permeabilized for 1 hour in 1% PBS-Triton, and blocked for 1 hour in WBR. Primary antibodies: mouse anti-Broad (1:500), rat anti-Ecadherin (1:50) (DSHB DCAD2), rabbit anti-Vasa (1:2000) (Paul Lasko) (Figure S2A), rat anti-Vasa (1:500) (Figure S3), mouse anti-Engrailed concentrate (1:20) or supernatant (1:50) (DSHB 4D9). Secondary antibodies: Alexa Fluor goat anti-mouse (1:200) and goat anti-rat (1:200). Ovaries were washed in PBT, incubated O/N at 4°C in PBT with primary antibodies and 5% NGS, washed in PBT, incubated for 1 hour at room temperature in secondary antibodies and DAPI, washed in PBT, mounted on slides and imaged on a Leica SP8X LSM.

Embryos - Females were aged for 2 days with wet yeast paste. ^~^100 males and ^~^100 females were mated and allowed to lay on apple juice plates O/N or for a few hours to obtain a range of desired stages. All aspects of this experiment prior to dechorionization were done at 25°C. Embryos were exposed to either 100% CO_2_ or air for 2 minutes by inserting the CO_2_ needle through a hole in the plastic fly bottle. The CO_2_ or air was humidified by flowing the gas through a bubbler and the flow rate was held precisely at 3.5 L per minute, using the Flow Buddy. Using a paint brush and embryo wash, embryos were washed from plates into a basket and rinsed with embryo wash using a squirt bottle, then dechorionated in 50% bleach (1:1 bleach:H_2_O) for 2 minutes by submerging the mesh end of the basket in a Falcon tube with the bleach. The de-chorionated embryos were rinsed with embryo wash in the basket, then transferred to a 1.5 mL Eppendorf tube by squirting embryo wash onto the outside end of the mesh and into the Eppendorf tube. Embryos were fixed for 40 minutes in 1 mL of heptane saturated with 37% formaldehyde. Using a pipette tip with the tip cut off and pre-treated in PBTA, embryos were transferred to Whatman paper for ^~^30 seconds to allow the heptane to evaporate. The Whatman paper was placed embryo side down on the tape, gently tapped until embryos were stuck to the tape, and the embryos were immediately covered with PBS. Vitelline membranes were manually removed using 30-Gauge syringe needles. Using a pipette controller attached to a glass Pasteur pipette pre-treated with PBTA, embryos were transferred to PBTA with 0.1% Triton X-100 for staining or to PBTA with 0.05% Triton X-100 for storage at 4°C. Stage 3-5 embryos were incubated with DAPI (1μg/mL), and rhodamine phalloidin (Abcam ab235138) (1:500) for 1 hour, and washed in PBTA with 0.05% Triton X-100 for 1 hour, rinsed 3x in PBS, and mounted in Vectashield mounting medium. Stage 8-12 embryos were incubated with rat anti-E-cadherin (1:50) and mouse anti-Engrailed concentrate (1:20) or supernatant (1:50) in PBTA with 0.1% Triton X-100 and NGS O/N at 4°C, washed for 1 hour in PBTA with 0.1% Triton X-100, incubated for 1 hour at RT in PBTA with 0.1% Triton X-100, 5% NGS, DAPI (1μg/mL), rhodamine phalloidin (Abcam ab235138) (1:500), Alexa Fluor goat anti-mouse 488 and anti-rat 647, washed for 1 hour in PBTA with 0.1% Triton X-100, rinsed 3x in PBS, and mounted on slides with Vectashield mounting medium.

#### Generation of *Idgf* mutant fly lines

Deletions of the entire locus or most of the locus for each of *Idgf1*, *Idgf2*, *Idgf3*, *Idgf4*, *Idgf5*, and *Idgf6* were created using CRISPR/Cas9-catalyzed homology-directed repair. For each deletion, we designed and injected three vectors: two guide RNAs and one donor containing a Ds-Red eye reporter to aid in screening newly transformed flies. Using the FlyCRISPR algorithm (tools.flycrispr.molbio.wisc.edu/targetFinder), we designed guide RNAs targeting sites upstream and downstream of each gene. Guide RNAs were synthesized as 5’-phosphorylated oligos (Operon), annealed, and ligated into the BbsI sites of the pU6-BbsI-chiRNA plasmid (Addgene #45946). For the donor vector, we PCR amplified ^~^1kb homology arms from genomic DNA of the injection strain and cloned them into pHD-DsRed-attP (Addgene #51019). We co-injected guide RNAs and donor vectors (Rainbow Transgenics) into *y vas-Cas9 w^1118^* embryos (Bloomington 55821) and screened F1 flies for germline transmission of the DsRed reporter, followed by single-fly PCR and sequencing. The *Idgf* null strains were outcrossed to *w^1118^; +; +* for three generations to remove background effects and remove the *vas-Cas9* transgene. All of the single *Idgf*-null lines are homozygous viable and fertile.

#### Image analysis and quantification

For egg chambers, images were acquired as 8-bit images at the same laser intensities and detector settings on a Leica SP8X LSM. Using ImageJ, we measured E-cadherin and Actin intensity across three cell membranes, from the center of one cell to the center of the neighboring cell, in at least 6 egg chambers for each genotype and compared egg chambers exposed to CO_2_ to egg chambers not exposed to CO_2_. Plot profiles across 5.0 μm were averaged over all egg chambers for each genotype and treatment.

For embryos, images of all genotypes and treatments were acquired as 16-bit images at the same laser intensities and detector settings on a Leica SP8X LSM. For each individual channel and optical plane being measured, background was subtracted, and a median filter was applied (rolling ball radius of 2.0) prior to measurements. For each genotype, we measured E-cadherin and Actin fluorescent intensity as above, across 6 cell membranes per embryo, with n=10 embryos for each treatment (no CO_2_, +CO_2_, and +air) within three bioreplicates, except for *sxtΔ* bioreplicate 3, no CO_2_ (n=6 embryos). Plot profiles across 3.5 μm were averaged over all embryos for each genotype and treatment. Total E-cadherin or Actin was calculated by integrating the area under the mean plot profile curve using the trapezoidal rule. Data analyses were performed in Excel. Statistical analyses were performed in Excel and R.

## Data Availability

Fly lines are available upon request. The authors affirm that all data necessary for confirming the conclusions of the article are present within the article, figures, tables, and in the supplementary data files: Data Files S1 through S8. Data file S1 contains hatch rate data, data file S2 contains dorsal appendage defect and rescue data, data file S3 contains hypoxia and CO_2_ exposure data, data file S4 contains Actin and E-cadherin profile data for egg chambers, data file S5 contains Actin and E-cadherin profile data in later stage embryos, data file S6 contains Actin and E-cadherin profile data in earlier stage embryos, data file S7 contains eclosion data, and data file S8 contains germ-cell count data.

## Acknowledgements

We thank the Bloomington *Drosophila* Stock Center (NIH P40OD018537) and the Vienna *Drosophila* Resource Center for stocks, and FlyBase for genetic, polypeptide, and functional data. The rabbit anti-Vasa antibody (Liu *et al*. 2009) was a gift from Paul Lasko. The mouse anti-Broad, rat anti-Vasa, and rat anti-E-cadherin antibodies were obtained from the Developmental Studies Hybridoma Bank, created by the National Institute of Child Health and Human Development of the National Institutes of Health (NIH) and maintained by the University of Iowa, Department of Biology. We thank Nathaniel Peters at the University of Washington Keck Center for technical support and advice on imaging. We thank Vincent So for providing the data and image for Idgf6::GFP localization to germ cells. We thank Dana Miller in the Department of Biochemistry at the University of Washington for technical assistance and advice on the hypoxia and CO_2_ experiments and Greg Beitel in the Department of Molecular Biosciences at Northwestern University for technical assistance and advice on the CO_2_ experiments.

## Author Contributions

Celeste Berg – Project design; oversight of all experiments; analysis of mutant embryos; review and editing of manuscript

Liesl Strand – Generation of *Idgf6Δ* fly strain using CRISPR/Cas9; experiments and data analysis to examine *Idgf6Δ* dorsal appendage defects; experiments to assess variability in phenotypes; review of manuscript.

Anne Sustar – Generation of *Idgf* null fly strains using CRISPR/Cas9; experiments (hatch rates, dorsal appendage morphogenesis, adult phenotype analysis, germ cell migration phenotype, CO_2_ response in egg chambers, patterning in egg chambers, Actin and E-cadherin measurements in egg chambers); imaging; data analyses; review of manuscript.

Sandra Zimmerman – writing and editing drafts, constructing figures, experiments (E-cadherin and Actin in embryos, hatch rates), imaging, and data analysis.

## Supplementary Figures and Tables

**Figure S1.**
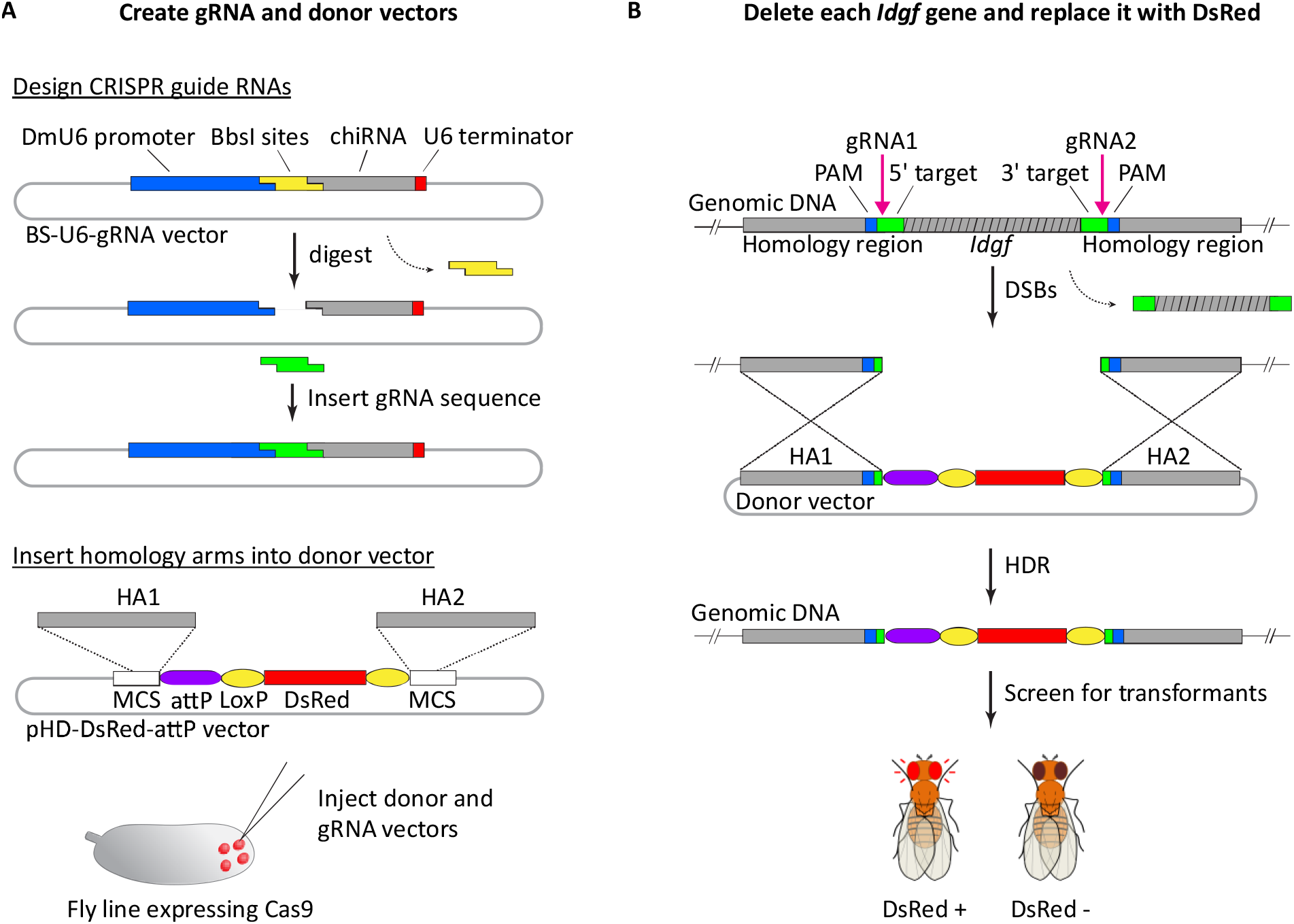
*Idgf* gene deletion method. (A) Strategy for constructing gRNA vectors and donor vectors for injection into embryos endogenously expressing Cas9. (B) Cas9 exploits gRNAs to make double-strand breaks at sites of pink arrows. Using homology directed repair, the *Idgf* gene is replaced with the gene encoding DsRed fluorescent protein. Transformants are null for the *Idgf* gene of interest and have fluorescent red eyes. gRNA (guide RNA), HA (homology arm), MCS (multiple cloning site) PAM (protospacer adjacent motif), DSB (double-strand break) HDR (homology directed repair).

**Figure S2.**
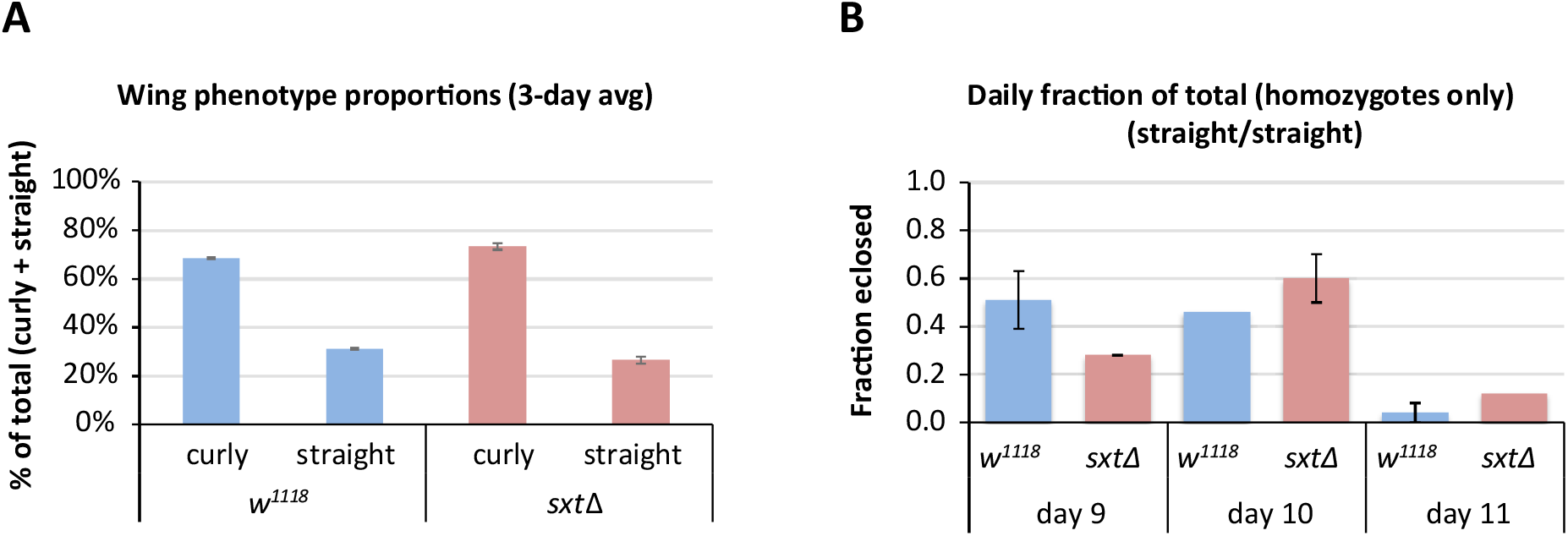
Eclosion rates. (A) Eclosion averaged over 3 days [9-11 days after egg laying (AEL)] for offspring of *+; +/CyO* and *w^1118^ Idgf^4Δ^*; *Idgf^(1-3)Δ,(6-5)Δ^/CyO* crosses. Percentages are the number of curly or straight-winged flies divided by the total (curly + straight). The expected Mendelian fractions of curly vs. straight-winged flies should be 2/3 heterozygous (curly-winged) and 1/3 homozyous (straight-winged) flies. The relative fractions of wing phenotypes in the *sxtΔ* background (73% curly, 27% straight) deviates significantly from the relative fractions in the *w^1118^* background (69% curly, 31% straight) (*p* = 0.0087, *X*^2^ test, n=1304 for the *w^1118^* background and 1185 for the *sxtΔ* background). (B) Eclosion rates for homozygous offspring over time. Fractions are the number of homozygous flies eclosed for each day divided by the total number of homozygous flies for all three days. The fraction of *w^1118^* homozygotes peaks 1 day earlier than for the *sxtΔ* homozygotes, indicating that the *sxtΔ* homozygotes are developmentally delayed.

**Figure S3.**
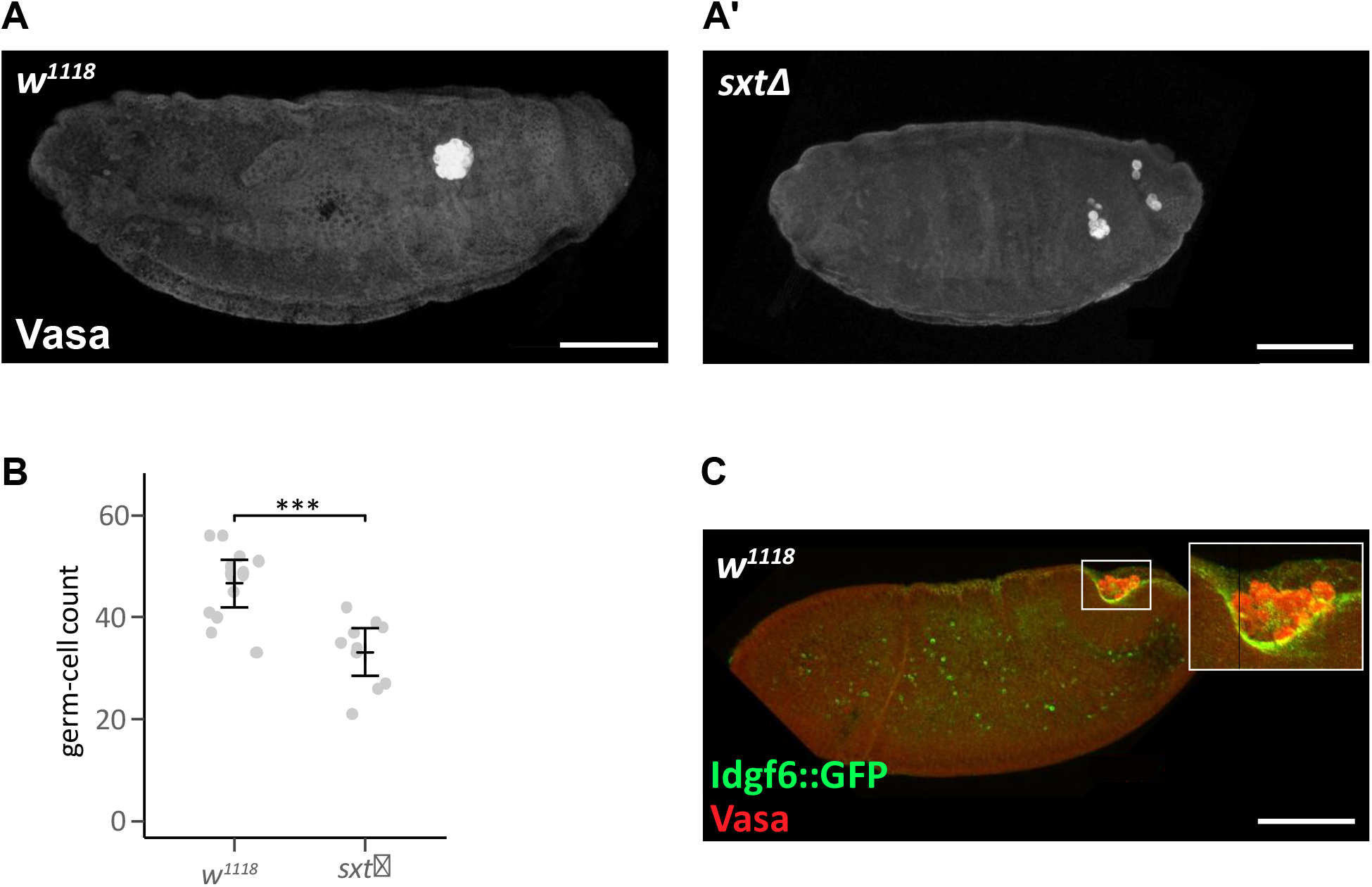
Germ cell migration. Anterior is to the left, dorsal is up in A, A’, and C. (A, A’) Germ cells in 16-hour old embryos are marked with anti-Vasa antibody, revealing germ-cell loss and reduced numbers in the gonads of sxtΔ versus *w^1118^* embryos. Scale bar = 50μm. (B) Graph shows germ cell counts in individual 3-hour old cellular blastoderm embryos. *w^1118^*, mean = 46.6 (n= 12 embryos), *sxtΔ*, mean = 33.2 (n= 10 embryos), *p* = 0.0002, Student’s *t*-test. (C) During germband extension, GFP-tagged Idgf6 (green) is present in and around the pole cells, which are marked with anti-Vasa antibody (red). Scale bars = 100μm.

**Figure S4.**
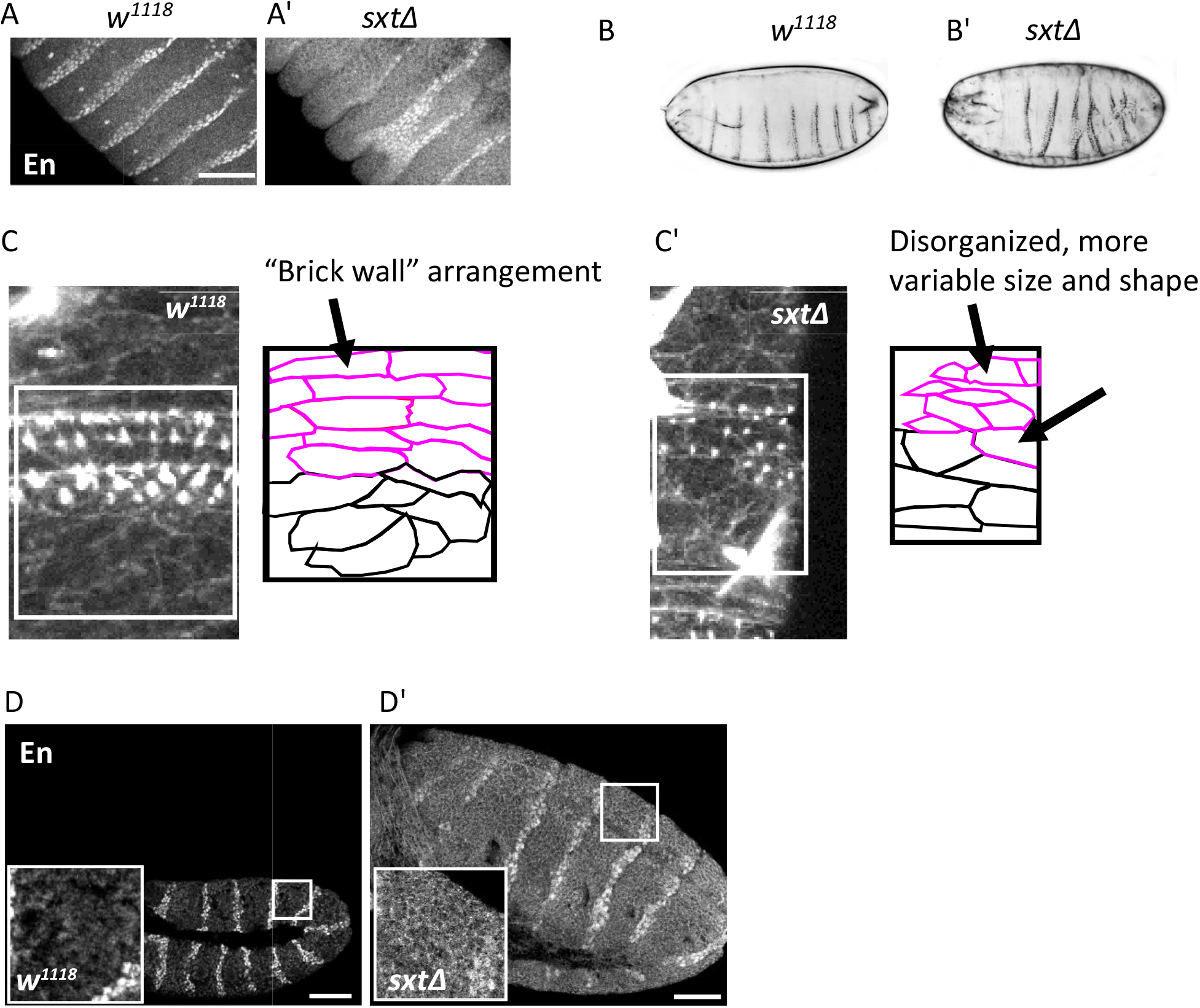
Segmentation defects. (A,A’) Anti-Engrailed antibody marks the posterior cells in each segment in *w^1118^* embryos. *sxtΔ* mutants exhibit segmentation defects. Scale bar = 50μm. (B,B’) Denticle belt defects in *sxtΔ* mutants reflect segmentation defects. Lateral view (B), ventrolateral view (B’). (C,C’) In Stage 16 embryos, Rhodamine-Phalloidin highlights cell cortices and the actin protrusions that secrete cuticle to form denticles (images). White squares in images show the areas drawn in the diagrams. (C) The control (*w^1118^*) cells display the typical “brick wall” arrangement in the denticle-producing cells (magenta) and the larger, less organized naked cuticle cells (black). (C’) Cell morphology and localization in denticle-producing cells is disrupted in *sxtΔ* mutant embryos. (D,D’) Engrailed protein aberrantly localizes to cell membranes in *sxtΔ* embryos (C’), but not in *w^1118^* embryos (C). White rectangle indicates area shown in close-up. Scale bar = 50μm.

**Figure S5.**
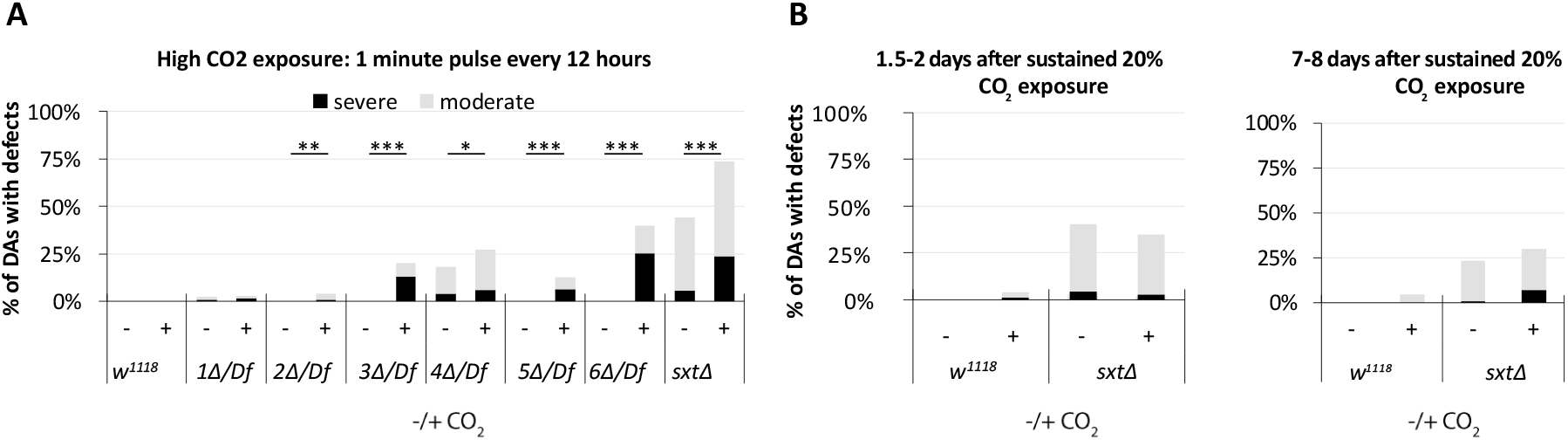
CO_2_ Exposure regimes. DA defects are significantly increased in *Idgf2Δ*, *Idgf3Δ*, *Idgf4Δ*, *Idgf5Δ*, *Idgf6Δ*, and *sxtΔ* mutants after a one-minute pulse of high (100%) CO_2_ exposure every 12 hours. A sustained time course of 20% CO_2_ for at least 1.5 days and as much as 8 days does not significantly increase DA defects in these mutants. * denotes *p*<0.05, ** denotes *p*<0.01, *** denotes p<0.001, Bonferroni adjusted *p*-values, *X*^2^ test for trends in binomial proportions (Rosner 2000).

**Figure S6.**
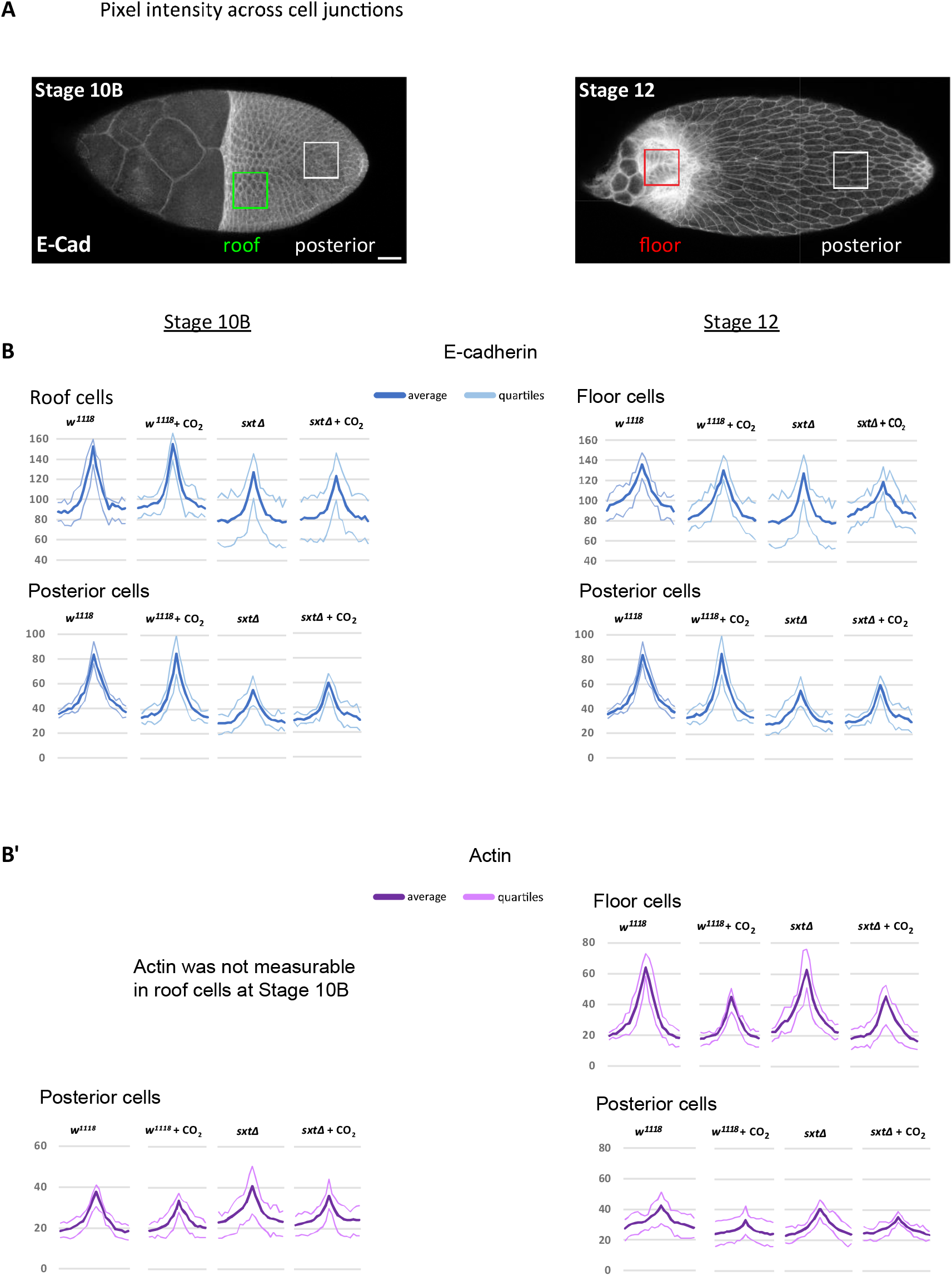
Quantification of cortical Actin and E-Cadherin intensity in DA cells. (A) E-Cadherin and Actin in Stage 10B roof cells (green square) and Stage 12 floor cells (red square) and in posterior columnar cells (white squares). (B) Comparison of *w^1118^* and *sxtΔ* fluorescence intensity plot profiles averaged across three cell membranes, from the center of one cell to the center of the neighboring cell, averaged over at least 6 egg chambers for each genotype and each exposure regime as indicated. Plots show mean (E-cad, blue and Actin, magenta) and confidence limits (gray).

**Table S1.** Molecularly mapped *Idgf* gene loci, mRNA, protein lengths and weights, and deleted sequences.

| Gene | Locus<sup>a</sup> | Length (bp) | Isoform | mRNA Length (nt) | Protein Length (aa) | MW (kDa) | Deleted Sequence<sup>b</sup> |
| --- | --- | --- | --- | --- | --- | --- | --- |
| *ldgf1* | 16,445,278 | 1516 | A | 1445 | 439 | 49.4 | 16,445,132 - 16,446,690 |
| *ldgf2* | 16,447,330 | 3179 | A | 1440 | 440 | 49.2 | 16,449,257 - 16,450,348 |
|  |  |  | B | 1796 | 368 | 41.1 |  |
|  |  |  | C | 1367 | 368 | 41.1 |  |
| *ldgf3* | 16,450,907 | 4288 | A | 1649 | 441 | 49.3 | 16,450,962 - 16,452,978 |
|  |  |  | B | 3145 | 441 | 49.3 |  |
|  |  |  | C | 1724 | 441 | 49.3 |  |
|  |  |  | D | 3801 | 441 | 49.3 |  |
| *ldgf4* | 9,927,973 | 2976 | A | 1514 | 442 | 48.6 | 9,928,084 - 9,930,904 |
|  |  |  | B | 1541 | 442 | 48.6 |  |
|  |  |  | C | 1807 | 442 | 48.6 |  |
| *ldgf5* | 18,393,666 | 1617 | A | 1495 | 444 | 50.1 | 18,393,347 - 18,395,294 |
| *ldgf6* | 16,691,022 | 2730 | A | 1541 | 452 | 50.4 | 16,690,912 - 16,693,893 |
|  |  |  | B | 1811 | 452 | 50.4 |  |

a. 5' nucleotide locus of molecularly mapped sequence from flybase.org
b. Cas9 cuts directly before the first and after the second number

**Table S2.**
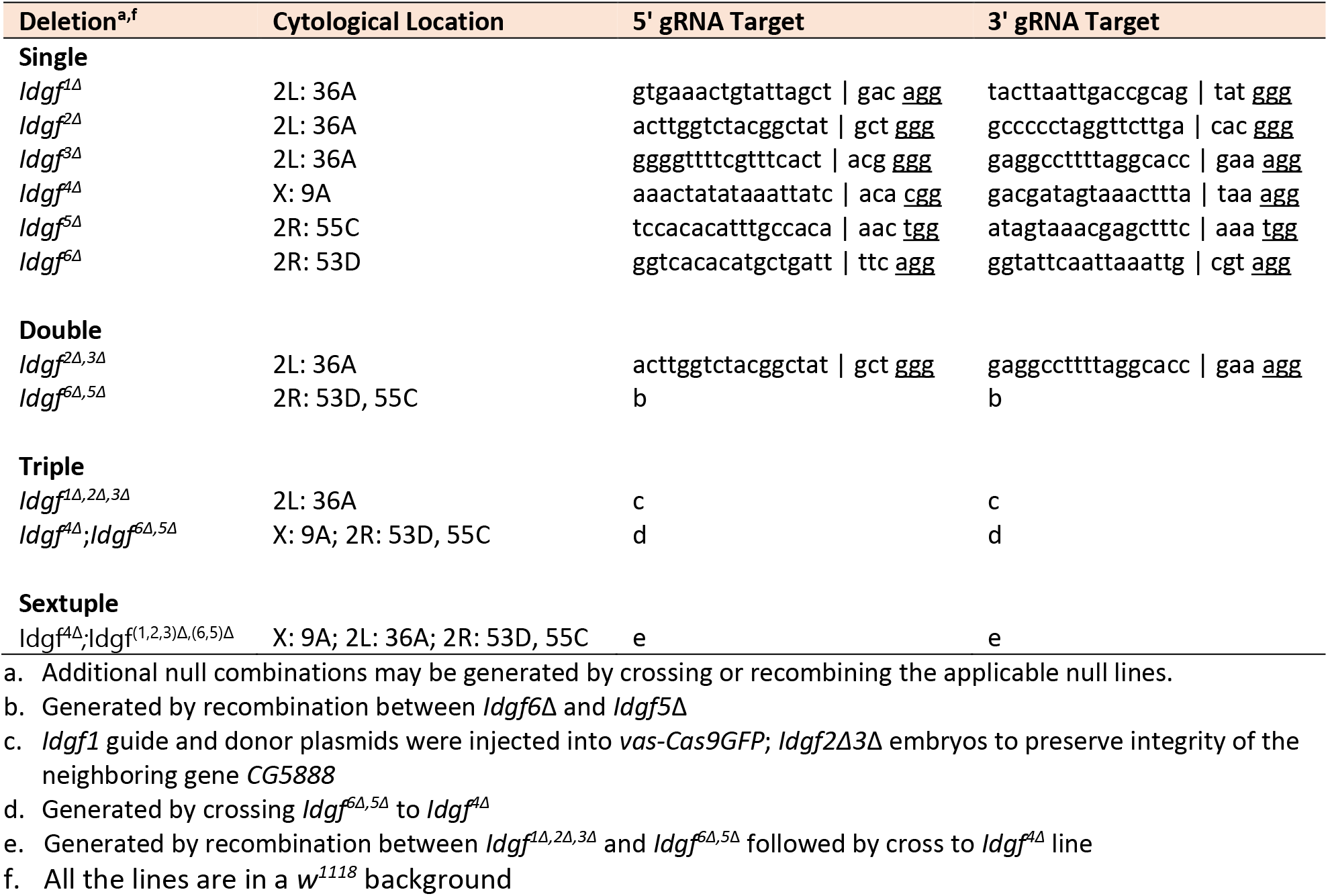
Cytological locations, gRNA sequences, and cut sites. Vertical line (|) indicates gRNA cut site, underlined nucleotides indicate PAM (located 3’ to target on genomic DNA, not part of gRNA)

**Table S3.** Deficiency phenotypes. The deficiencies delete the indicated Idgfs. Deficiency locus is heterozygous with wild-type Idgf(s) to test for DA defects caused by deletion of genes flanking the Idgfs within the deletion locus.

| genotype | associated Idgf deletions | normal | moderate | severe |
| --- | --- | --- | --- | --- |
| $w^{1118}; +/+$ | NA | 98.4% | 1.5% | 0.1% |
| $w^{1118}; +/Df(2L)ED1102$ (VENKEN *et al.* 2006); + | 1Δ, 2Δ, 3Δ | 99.1% | 0.7% | 0.1% |
| $w^{1118}/Df(1)ED6989$ (VENKEN *et al.* 2006) | 4Δ | 100.0% | 0.0% | 0.0% |
| $w^{1118}; +/Df(2R)BSC337; +$ | 5Δ | 99.8% | 0.2% | 0.0% |
| $w^{1118}; +/Df(2R)6064; +$ | 6Δ | 99.4% | 2.0% | 0.0% |

